# Cerebellar tonic inhibition orchestrates the maturation of information processing and motor coordination

**DOI:** 10.1101/2024.05.30.596563

**Authors:** Jea Kwon, Sunpil Kim, Junsung Woo, Keiko Tanaka-Yamamoto, Oliver James, Erik De Schutter, Sungho Hong, C. Justin Lee

## Abstract

Tonic inhibition in cerebellar granule cells (GC) is crucial in information coding fidelity for motor coordination. It emerges in activity-dependent and-independent manners, and their interplay evolves with age. However, specific molecular and cellular mechanisms and how their change affects network-level computation and motor behavior remain unclear. Here we show that, while net tonic inhibitory current remains unchanged, the main source of tonic GABA switches from synaptic spillover (neuronal activity-dependent) to astrocytic Best1 (activity-independent) throughout adolescence (4-8 weeks) in mice. Computational modeling based on experimental data demonstrated that this switch down-regulates the internally generated network activity mediating mutual inhibition between GC clusters receiving different inputs, thereby enhancing their independence. Consistent with simulations, 3D-posture analysis revealed an age-dependent increase in independent limb movements during spontaneous motion, which was impaired in Best1 knockout mice. Our findings highlight the late-stage development of complex motor coordination driven by the emergence of astrocyte-mediated tonic inhibition.

- 6 main figures, 58 references, 5,924 words in the manuscript (excluding the abstract, tables, figure legends and references).
- Supplementary information: 1 text (supplementary methods), 9 figures, 4 movies, and 1 table.

## Introduction

GABA (*γ*-aminobutyric acid), a primary inhibitory neurotransmitter, has two distinct modes of action: phasic and tonic inhibition^1–3^. Phasic inhibition is triggered by synaptic input and is characterized by being transient and rapidly desensitizing. In contrast, tonic inhibition relies on extrasynaptic GABA and provides a persistent conductance^4,5^ although receptor desensitization can still occur^6^. Multiple mechanisms have been identified in regulating extrasynaptic GABA levels, including spillover of synaptic GABA, uptake by GABA transporters (GATs), and Bestrophin-1 (Best1) channel-mediated GABA release from astrocytes^5,7^.

In particular, tonic inhibition affects cerebellar granule cells (GC), the most abundant neuronal cell type in the brain. Tonic inhibitory conductance in GCs contributes to the intrinsic membrane conductance and is, therefore, a crucial regulator of excitability, demonstrated *ex vivo*^8,9^ and *in vivo*^10^. Studies have suggested that GC tonic inhibition can regulate the signal-to-noise ratio of GC spike coding^8–11^ and modulate GC output in additive^8^ or multiplicative manners^12,13^.

A crucial but less well-known aspect of GC tonic inhibition is the developmental transition in its origin. In younger animals, the primary contribution comes from neuronal activity-dependent spillover GABA^14,15^. On the other hand, in adults, the main source of tonic inhibition is neuronal activity-independent ambient GABA^9,16^, most likely due to astrocytic GABA release mediated by Best1^17,18^. However, the functional implications of this developmental transition and the role of astrocytic GABA remain unclear. One potential implication of the age-dependent GC tonic inhibition can be late-stage sensorimotor learning. In humans, some cerebellum-related sensorimotor behavior begins to manifest only around the onset of adolescence^19,20^, which is paradoxically well beyond the currently known period for maturation of the cerebellar neurons and circuits. This raises questions: How does the developmental transition in the cellular source of tonic inhibition occur, even after the apparent maturation of the cerebellar neural circuit? And how does it impact network computation and behavior?

This study investigates these questions by combining *ex vivo* experiments that systemically analyze the primary origin of tonic inhibition, *in silico* network modeling based on experimental data, and *in vivo* behavioral analysis for both young (3-to 4-week-old) and adult (8-to 12-week-old) mice. Briefly, we performed a step-by-step elimination of the GABA spillover, GATs, Best1, and their combinations in *ex vivo* experiments to measure their contribution to tonic inhibition in both groups, which revealed that, despite the same level of tonic inhibition, its primary source is GABA spillover from synaptic activity during youth while it is predominantly astrocytic Best1 channels in adulthood. The increased uptake of spillover GABA by GATs is also found to play a pivotal role in this transition. The large-scale computational models integrated with experimental data demonstrated that the shift from neuronal activity-dependent toward independent inhibition decreases the network-wide regulation of the GC clusters activated by distinct mossy fiber inputs, promoting their independence and potentially allowing similar phenomena in sensorimotor control. Finally, our analysis of animal posture during spontaneous movements showed that animals exhibited increased movement diversity and flexibility, involving more independent limb movements as they mature, which was disrupted in Best1 knockout mice.

## Materials and Methods

### Animals

Young (3-to 4 weeks old) and adult (8-to 12 weeks old) WT and Bestrohpin-1 knockout male mice (BALB/c background) were used for both *ex vivo* slice patch clamping and behavior experiments. Mice were given *ad libitum* access to food and water under a 12:12-h light-dark cycle. Animals were housed in groups of 3–5 per cage. All experimental procedures were conducted according to protocols approved by the directives of the Institutional Animal Care and Use Committee of IBS (Daejeon, Republic of Korea).

### Preparation of brain slices

Mice were deeply anesthetized using vaporized isoflurane and then decapitated. After decapitation, the brain was quickly excised from the skull and submerged in ice-cold, oxygenated (95% O_2_/5% CO_2_) sucrose-based cutting solution that contained 5 KCl, 1.23 NaH_2_PO_4_, 26 NaHCO_3_, 10 glucose, 0.5 CaCl_2_, 10 MgSO_4_, and 212.5 sucrose (in mM) with pH 7.4. The hemisected brain was glued onto the stage of a vibrating microtome (PRO7N; DSK) and 300-*µ*m-thick sagittal cerebellar slices were cut and transferred to an extracellular aCSF solution containing 130 NaCl, 24 NaHCO_3_, 3.5 KCl, 1.25 NaH_2_PO_4_, 1.5 CaCl_2_, 1.5 MgCl_2_ and 10 D-(+)-glucose (in mM) with pH 7.4. Slices were incubated at room temperature for at least 1 h before recording, with oxygenation (95% O_2_/5% CO_2_).

### Cell-attached recording

To measure E_THIP_, we measured DF_THIP_ and V_m_ as previously described ^21–24^. For both of the experiments, we applied fire polishing followed by the Sylgard coating to enhance the seal and precision of single-channel recordings. For the high-KCl internal solution, the pipette was filled with an internal solution consisting of 120 KCl, 35 KOH, 1 CaCl_2_, 10 HEPES, 11 EGTA, 2 MgCl (in mM) with pH 7.3. To establish the reversal potential of potassium currents and activate voltage-gated potassium channels, depolarizing voltage ramps ranging from a holding potential of-100 mV to +200 mV were applied. During the intervals between stimulations, the patch was held to-60 mV with respect to the membrane potential (Vm) to mitigate potential voltage-dependent “steady-state” inactivation effects from the K(V) channel at the physiological V_m_. For measuring the DF_THIP_, the pipette was filled with an internal solution consisting of 120 NaCl, 5 KCl, 0.1 CaCl_2_, 10 MgCl_2_, 20 TEA-Cl, 5 4-AP, 10 HEPES, 10 glucose (in mM) with pH 7.35 and 100 *µ*M of THIP was freshly added before the experiment.

DF_THIP_ was determined as the potential where single-channel currents reversed polarity. Liquid junction potentials were not corrected and would introduce maximal shifts of ±2.1 mV.

### Tonic GABA_A_R current recording

Slices were transferred to a recording chamber that was continuously perfused with oxygenated (95% O_2_/5% CO_2_) aCSF solution. The slice chamber was mounted on the stage of an upright microscope and viewed with a 60× water immersion objective (numerical aperture = 0.90) with infrared differential interference contrast optics. Cellular morphology was visualized by a complementary metal oxide semiconductor camera and the Imaging Workbench software (INDEC BioSystems, ver. 9.0.4.0). Whole-cell recordings were made from granule cells located in the cerebellum. The holding potential was-60 mV. Pipette resistance was typically 6–8 MΩ and the pipette was filled with an internal solution consisting of 135 CsCl, 4 NaCl, 0.5 CaCl_2_, 10 HEPES, 5 EGTA, 2 Mg-ATP, 0.5 Na_2_-GTP, and 10 QX-314 (in mM) with pH 7.2 adjusted by CsOH (278–285 mOsmol). Before measuring the tonic current, the baseline current was stabilized with D-AP5 (50 *µ*M) and CNQX (20 *µ*M). For the blockade of action potential-dependent spillover, the voltage-gated sodium channel was blocked with TTX (1 *µ*M). For the blockade of GABA uptake, GABA transporters were blocked with a pan-GAT inhibitor, Nipecotic acid (1 mM). Electrical signals were digitized and sampled at 10 ms intervals with the Digidata 1550 data acquisition system and the Multiclamp 700B Amplifier (Molecular Devices) using the pClamp10.2 software. Data were filtered at 2 kHz. The amplitude of the tonic GABA current was measured by the baseline shift after gabazine (GBZ) (50 *µ*M) administration using the Clampfit software (ver. 10.6.0.13.). The frequency and amplitude of spontaneous inhibitory postsynaptic currents before bicuculline administration were detected and measured by Mini Analysis (Synaptosoft, version 6.0.7.).

### *In silico* model of the granular layer network in the cerebellar cortex

We built and simulated the physiologically detailed large-scale computational model of the cerebellar granular layer ^25,26^ with parameters tuned to our experimental data. Briefly, the model corresponded to a 1.5 mm × 0.7 mm region of the granular layer in the cerebellar cortex, containing about 0.8 million GCs, 2000 GoCs, and 2000 MFs. We determined the number and properties of GABAergic synapses and tonic Cl^-^ conductance in GCs by comparing experimental data and simulations in the cases of WT adult, WT young, Best1 KO adult, and Best1 KO young animals. After the parameter tuning, the network models were constructed and simulated via the previously described procedures ^25,26^ on the OIST supercomputer Deigo (https://groups.oist.jp/scs/deigo). The models will be available via modelDB (https://modeldb.yale.edu). For more details, see Supplementary Methods.

### Capturing and analysis of animal behavior

Key body points (marker positions) of mice were extracted by the AVATAR system as previously described^27,28^. Briefly, mouse movement in an open-field chamber (transparent cuboid, 20 cm (w) × 20 cm (l) x 30 cm (h)) was recorded over a period of 10 minutes using five cameras positioned on four sides and one at the bottom, operating at a frequency of 20 Hz. Nine key body points of the subjects — including the nose, neck, chest, anus, tail tip, left forepaw, right forepaw, left hindpaw, and right hindpaw — were identified and generated through the YOLO-based DARKNET model, followed by a detailed 3D reconstruction algorithm.

From the extracted marker positions (nodes), we computed variables to characterize the animal movement. In particular, we computed the angle between the limb and body vectors^29,30^ (Fig. 6b) to characterize limb coordination by their correlations during periods of large movement. The large movement was selected by the criterion that the velocity of the anus node exceeds 2 cm/s, and the fore-to-hind paw vertical distance is within the upper limit. The limit was set by 2.576 times the standard deviation of the vertical distance between the left and right hindpaws, imposing that the forepaws are not higher than the maximum height that the hindpaws can reach 99% of the time. We also segmented and clustered the marker position time series by analyzing movement features and computed the angle-angle correlations per cluster and how they depend on the body rotation speed (Fig. 6f-g). For more details, see Supplementary methods.

### Immunohistochemistry

Primary antibodies used were rabbit anti-GABRα1 (06-868, Sigma), rabbit anti-VGAT (131 003, Synaptic Systems), and mouse anti-gephyrin (147 111, Synaptic Systems). Secondary antibodies were Alexa Fluor 488-conjugated anti-rabbit and Alexa Fluor 555-conjugated anti-mouse IgG antibodies (Thermo Fisher Scientific).

All procedures involving mice were conducted in accordance with the guidelines of the Institutional Animal Care and Use Committee of Korea Institute of Science and Technology. For IHC analysis, male C57BL6J mice were used (n=8 adult, 8–10 weeks old; n = 8 young, 3–4 weeks old). Mice were anesthetized with isoflurane and perfused transcardially with glyoxal fixative^31^ (9% glyoxal, 8% acetic acid, pH 7.4). Brains were immediately removed, post-fixed overnight in glyoxal fixative at 4°C, and sectioned sagittally into 50-µm cerebellar slices using a vibratome (Leica VT1200S). Slices were blocked for 30 min at room temperature in 5% normal goat serum in PBS, incubated with primary antibodies overnight at 4°C, washed several times, and incubated with secondary antibodies overnight at 4°C. Finally, slices were mounted and imaged using an AX laser-scanning confocal microscope (Nikon) with low (10×, 1,024 × 1,024 resolution) and high (60×, 2,048 × 2,048 resolution) magnification objectives.

Image analyses were performed using NIS-Element (Nikon), ImageJ (National institute of Health, https://imagej.net/ij/), and Fiji^32^ (https://imagej.net/software/fiji/). In low-magnification images, regions corresponding to the granular and molecular layer of lobule IV/V were manually selected, and signal intensities were measured. For quantification, signal intensities in the GL were normalized to those in the ML.

Statistical differences were assessed using the Mann–Whitney test in OriginPro software. Quantitative IHC data are presented as boxplots, with black dots indicating individual data points, center lines denoting the median, open squares representing mean values, boxes spanning the 25th to 75th percentiles, and whiskers indicating the minimum and maximum values.

### Statistical analysis

The numbers and individual dots refer to the number of brain slices or animals unless otherwise clarified in the figure legends. For data presentation and statistical analysis, GraphPad Prism 9.1.2 (GraphPad Software) and Python were used. For comparing two groups, the Mann-Whitney U test or Student’s t-test was used. For comparing more than two groups, one-way ANOVA with Tukey’s multiple comparisons test, Kruskal-Wallis test with Dunn’s multiple comparisons test, or two-way ANOVA with Sidak’s multiple comparisons test was used. Statistical significance was set at *p < 0.05, **p < 0.01, ***p < 0.001, ****p < 0.0001. For error bars in Fig. 5f-g and 6f-g, we employed resampling (1000 iterations) to generate distributions, from which we computed and used the standard error of the mean (SEM). All data are presented as mean ± SEM unless otherwise stated.

## Results

### Differential modulation of ambient GABA level in the young and adult cerebellum

We explored age-related changes in ambient GABA levels in the cerebellum by performing whole-cell patch clamp recording on GCs and measuring the tonic GABA_A_ currents. Our first focus was on action potential-dependent GABA spillover that was found to be one of the major GABA sources previously (Fig. 1a). To identify age-related changes in the spillover effect on extrasynaptic GABA level, we sequentially applied tetrodotoxin (TTX), an inhibitor of action potential-mediating voltage-gated sodium channels, and gabazine (GBZ), a GABA_A_R antagonist (Fig. 1b, d). TTX treatment positively shifted the holding currents in the young but not in the adult condition, whereas GBZ treatment affected both conditions (Fig. 1c, e). In adult GC, the TTX-sensitive component was smaller, whereas the TTX-insensitive component was larger compared to young GCs despite no significant difference in the total amount of tonic current (Fig. 1h): the TTX-sensitive fraction of the tonic current was 63.5% and 22.6% in young and adult, respectively (Fig. 1j).

**Figure 1.**
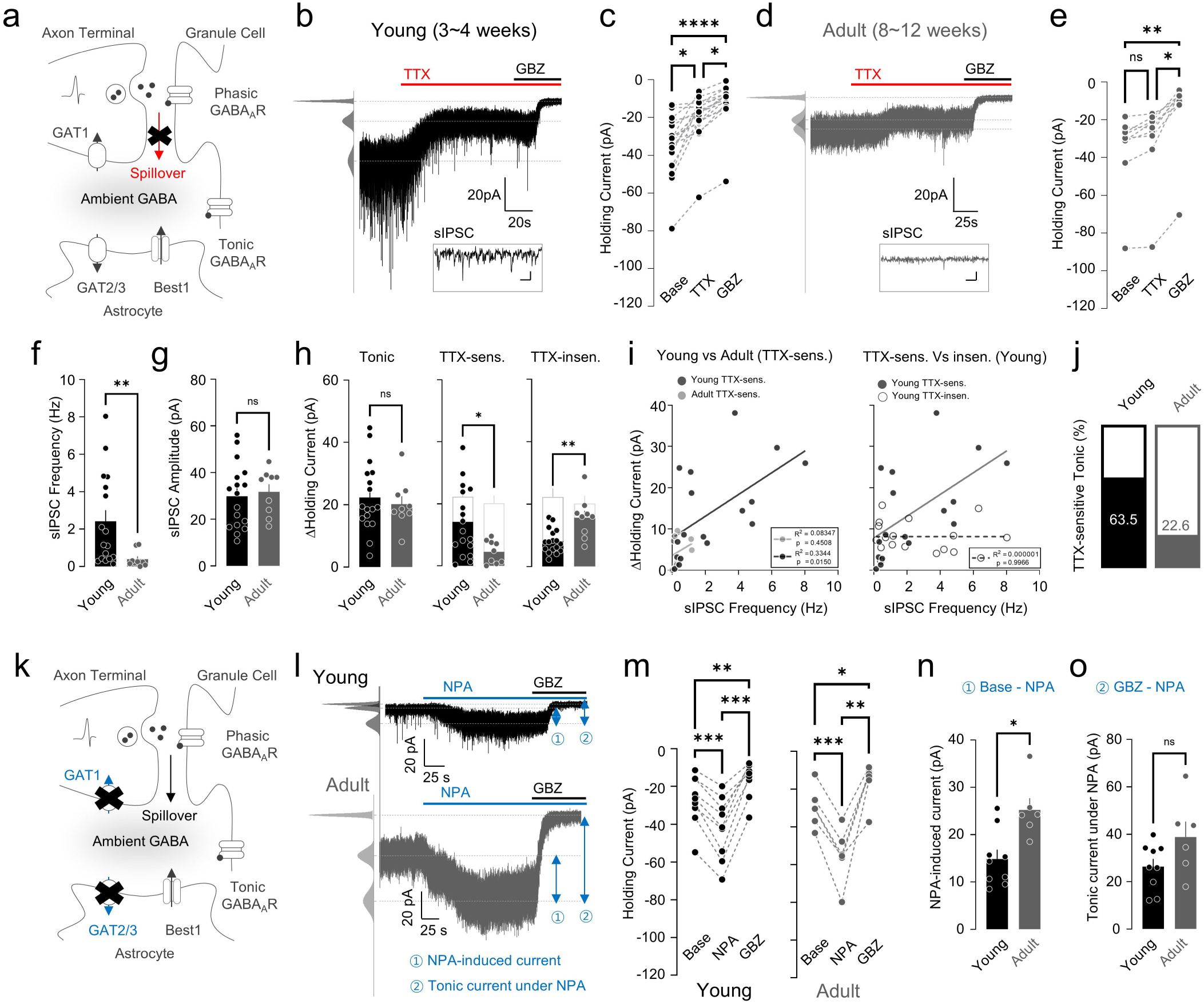
Spillover-and GAT-mediated tonic current in young and adult mice. **a,** Model of spillover-mediated (red) ambient GABA. **b,** Representative trace of GC tonic current from a young mouse applied with TTX and GBZ; inset, enlarged sIPSC. **c,** Paired scatter plots of holding current change. Kruskal-Wallis test (****p < 0.0001), Dunn’s multiple comparisons test, Base vs TTX, *p = 0.0107; TTX vs GBZ, *p = 0.0107; Base vs GBZ, ****p < 0.0001. **d,** Representative trace of tonic current from GC of adult mouse applied with TTX and GBZ; inset, enlarged sIPSC. **e,** Paired scatter plot of holding current change. Kruskal-Wallis test (****p < 0.0001), Dunn’s multiple comparisons test, Base vs TTX, p = 0.9999; TTX vs GBZ, *p = 0.0411; Base vs GBZ, **p = 0.0026. **f,** Summary scatter bar graph of sIPSC Frequency. Mann-Whitney test, Young vs Adult, **p = 0.0029. **g,** Summary scatter bar graph of sIPSC Amplitude. Unpaired Student’s t-test, Young vs Adult, p = 0.7171. **h,** Summary scatter bar graphs of tonic current (left), TTX-sensitive current (middle), and TTX-insensitive current (right). Unpaired Student’s t-test, Tonic: Young vs Adult, p = 0.6134; TTX-insensitive: Young vs Adult, *p = 0.0196; TTX-sensitive: Young vs Adult, **p = 0.001. **i,** Linear regression analysis of sIPSC Frequency and ΔHolding current. Adult TTX-sensitive current vs Young TTX-sensitive current (left). Young TTX-sensitive current vs young TTX-insensitive current (right). Simple linear regression, Adult TTX-sensitive (R^2^ = 0.08347, p = 0.4508); Young TTX-sensitive (R^2^ = 0.3344, p = 0.0150); Young TTX-insensitive (R^2^ = 0.000001, p = 0.9966). **j,** TTX-sensitive portion of tonic current in young and adult mice. **k,** Graphical description of GABA transporter-mediated ambient GABA model. **l,** Representative traces of tonic current from GCs of young (top) and adult (bottom) mouse applied with NPA and GBZ. **m,** Paired scatter plots of holding current change in young (left) and adult (right). Young: RM one-way ANOVA (****p < 0.0001), Tukey’s multiple comparisons test, Base vs NPA, ***p = 0.0002; Base vs GBZ, **p = 0.0016; NPA vs GBZ, ***p = 0.0001; Adult: RM one-way ANOVA (****p < 0.0001), Holm-Sidak’s multiple comparisons test, Base vs NPA, ****p = 0.0005; Base vs GBZ, *p = 0.0259; NPA vs GBZ, **p = 0.0039). **n,** NPA-induced current in young and adult mice. Mann Whitney test, Base – NPA: Young vs Adult, *p = 0.0120. **o,** Tonic current under NPA in young and adult mice. Mann Whitney test, GBZ – NPA: Young vs Adult, p = 0.0879). GBZ: gabazine, GABA_A_ receptor blocker; TTX: tetrodotoxin voltage-gated sodium channel blocker; NPA: nipecotic acid, pan GABA transporter blocker.

We questioned whether this age-dependent decrease in the TTX-sensitive component is due to 1) a decrease in synaptic inputs and/or 2) an increase in GABA uptake from GABAergic axon terminals. First, we found a significant difference in spontaneous inhibitory post-synaptic current (sIPSC) frequency without changing amplitude between young and adult groups (Fig. 1f, g), indicating that presynaptic GABAergic inputs decrease in GCs with maturation. In addition, linear regression analysis revealed that sIPSC frequency has a significant correlation with young TTX-sensitive components but not with adult TTX-sensitive components or young TTX-insensitive components (Fig. 1i). This result supports that the reduction of TTX-sensitive tonic current is due to a decrease in synaptic input. We also checked the second possibility by applying NPA, a pan GAT inhibitor, before GBZ treatment and monitoring the contribution of GABA uptake in ambient GABA modulation (Fig. 1k, l). Both young and adult GCs exhibited NPA-induced tonic GABA_A_R current (Fig. 1m), but it was significantly larger in the adult conditions (Fig. 1n), suggesting an age-dependent increase in capacity for uptake of extrasynaptic GABA. Taken together, these results show that the TTX-sensitive component in tonic inhibition declines in an age-dependent manner due to a decrease of action-potential dependent GABA spillover (Fig. 1j) and an increase in GABA uptake (Fig. 1o).

### GATs-mediated GABA uptake affects action potential-dependent and independent tonic GABA differentially with age

Next, we estimated the age-dependent changes of TTX-sensitive and TTX-insensitive GABA release (Fig. 2a) in the absence of GABA uptake by recording tonic GABA_A_ currents in the presence of NPA (Fig. 2b). We found that the amount of net GABA release was not significantly different between young and adult mice (Fig. 2d). Notably, a previously unseen TTX-sensitive component (Base vs. TTX, ns; Fig. 1e) appeared under blockade of GATs in adults (Fig. 2c), indicating that most of the action potential-dependent GABA spillover was taken up by GATs in adults. Consistently, the TTX-sensitive component was not significantly different between young and adult animals under NPA treatment (Fig. 2d), suggesting that the amount of action potential-dependent GABA spillover is comparable between the two age groups. NPA treatment also significantly increased the TTX-insensitive component only in adults (Fig. 2e), implying that uptake of action potential-independent GABA release is differentially modulated by GATs between young and adult mice. Taken together, the relative contribution of GATs on the action potential-dependent and independent tonic GABA changes with aging, with an increased contribution of GATs to the TTX-sensitive GABA release in adults (Fig. 2f).

**Figure 2.**
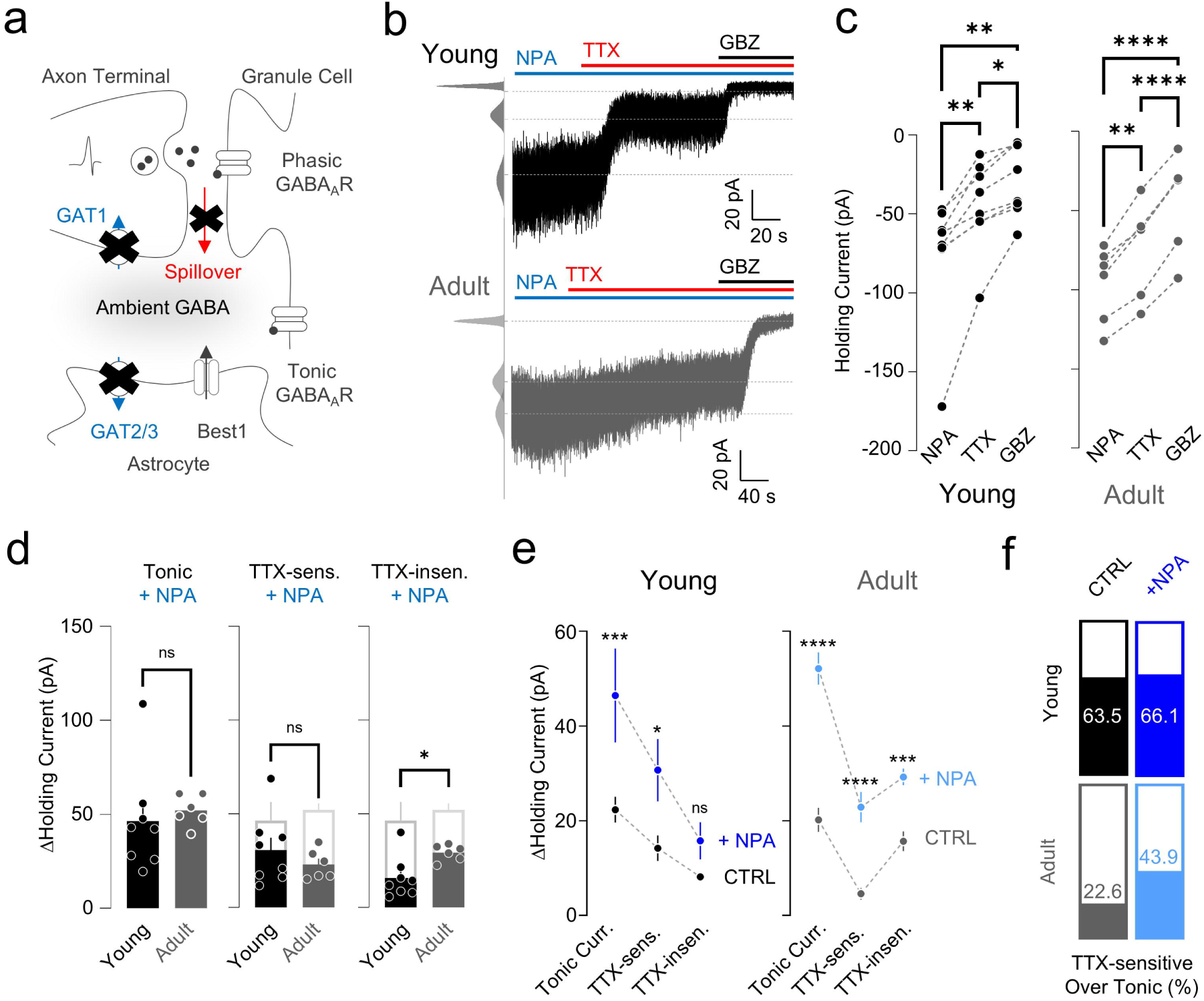
TTX-sensitive and TTX-insensitive tonic currents under blockade of GATs. **a,** Model of GAT-(blue) and spillover-mediated (red) ambient GABA. **b,** Representative traces of GC tonic current from a young (top) and adult (bottom) mouse applied with TTX and GBZ under NPA treatment. **c,** Paired scatter plots of holding current change in young (left) and adult (right). Young: RM one-way ANOVA (**, p=0.0018), Tukey’s multiple comparisons test, NPA vs TTX, **p = 0.0057; NPA vs GBZ, **p = 0.0055; TTX vs GBZ, *p = 0.0119; Adult: RM one-way ANOVA (****, p<0.0001), Tukey’s multiple comparisons test, NPA vs TTX, **p = 0.0020; NPA vs GBZ, ****p < 0.0001; TTX vs GBZ(****, p<0.0001). **d,** Summary scatter bar graphs of tonic current (left), TTX-sensitive current (middle), and TTX-insensitive current (right). Mann Whitney test, Tonic: Young vs Adult, p = 0.1812; TTX-sensitive: Young vs Adult, p = 0.4908; TTX-insensitive: Young vs Adult, *p = 0.0200. **e,** NPA effect on ΔHolding current in young (left) and adult (right) mice. Young: Two-way ANOVA, Sidak’s multiple comparisons test, Tonic: WT vs +NPA, ***p = 0.0003; TTX-sensitive: WT vs +NPA, *p = 0.0194; TTX-insensitive: WT vs +NPA, p = 0.4884; Adult: Two-way ANOVA, Sidak’s multiple comparisons test, Tonic: WT vs +NPA, ****p < 0.0001; TTX-sensitive: WT vs +NPA, ****p < 0.0001; TTX-insensitive: WT vs +NPA, ***p = 0.0009. **f,** TTX-sensitive tonic current portion in young (top) and adult (bottom) mice. GBZ: gabazine, GABAa receptor blocker; TTX: tetrodotoxin voltage-gated sodium channel blocker; NPA: nipecotic acid, pan GABA transporter blocker. NPA-containing solution was perfused from the beginning of the patching process onto the slice, which effectively resulted in ∼5–10 minutes of pre-incubation by the time the cell was patched.

### Best1 channel mediates action potential-independent GABA release

The non-synaptic GABA release has been associated not only with the Best1-mediated tonic release from glia but also with a reverse mode of GATs^7,33^. However, in our experiments, the primary role of GATs was uptake but not release (Fig. 2e). Thus, we hypothesized that Best1 mediates the remaining TTX-insensitive, non-synaptic component in both young and adult conditions (Fig. 3a). We tested this idea by comparing tonic currents from Best1 knockout (KO) with Best1 wild type (WT) cerebellar slices in both young and adult conditions (Fig. 3b, c). We observed a significantly lower tonic current in Best1 KO than WT in both young and adult mice (Fig. 3e), indicating that Best1 accounts for more than half of tonic currents in both age groups. Especially in adults, the reduced tonic current was primarily due to a significant decrease in non-synaptic components (Fig. 3e). As expected, the TTX-sensitive GABA release was not altered in both young and adult Best1 KO compared to WT (Fig. 3f). Taken together, these results imply that the Best1 channel makes a significant contribution to the action potential-independent tonic inhibition in both young and adult mice.

**Figure 3.**
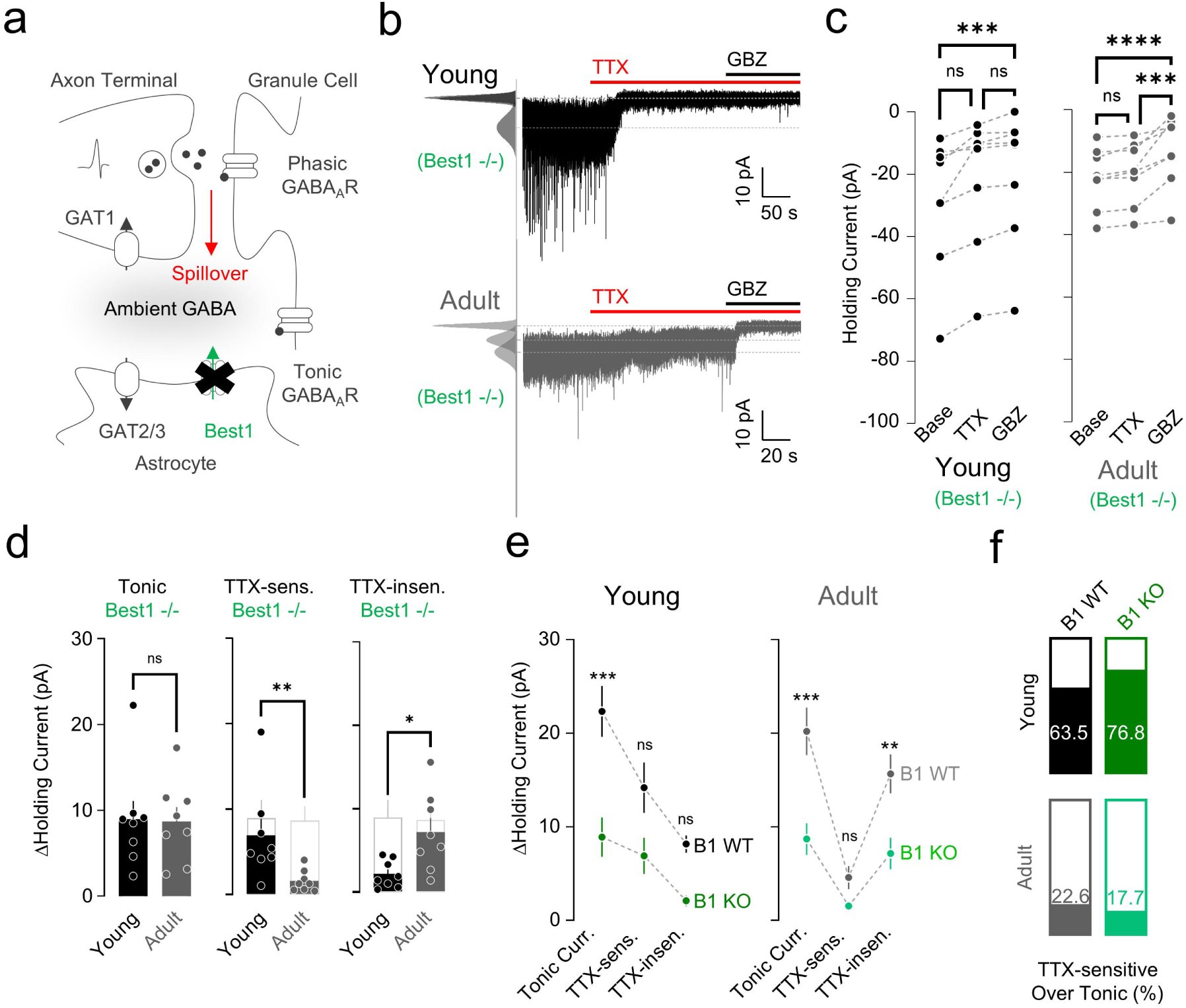
TTX-sensitive and TTX-insensitive tonic currents in the absence of Best1. **a,** Model of Best1-(green) and spillover-mediated (red) ambient GABA. **b,** Representative traces of GC tonic current from a young (top) and adult (bottom) mouse applied with TTX and GBZ in Best1-/-mice. **c,** Paired scatter plots of holding current change in young (left) and adult (right). Young: Friedman test (****, p<0.0001), Dunn’s multiple comparisons test, Base vs TTX, p = 0.1365; Base vs GBZ, ***p = 0.0002; TTX vs GBZ, p = 0.1365; Adult: RM one-way ANOVA (****, p<0.0001), Tukey’s multiple comparisons test, Base vs TTX, p = 0.5253; Base vs GBZ, ****p < 0.0001; TTX vs GBZ, ***p = 0.0004. **d,** Summary scatter bar graphs of tonic current (left), TTX-sensitive current (middle), and TTX-insensitive current (right). Unpaired Student’s t-test, Tonic: Young vs Adult, p = 0.9140; Mann Whitney test, TTX-sensitive: Young vs Adult, **p = 0.0030; Unpaired Student’s t-test, TTX-insensitive: Young vs Adult, *p = 0.0124). **e,** Best1 effect on ΔHolding current in young (left) and adult (right) mice. Young: Two-way ANOVA, Sidak’s multiple comparisons test, WT vs B1 KO, ***p = 0.0009; TTX-sensitive: WT vs B1 KO, p = 0.1250; TTX-insensitive: WT vs B1 KO, p = 0.2476; Adult: Two-way ANOVA, Sidak’s multiple comparisons test, Tonic: WT vs B1 KO, ***p = 0.0001; TTX-sensitive: WT vs B1 KO, p = 0.5526; TTX-insensitive: WT vs B1 KO, **p = 0.0045. **f,** TTX-sensitive tonic current portion in young (top) and adult (bottom) mice. GBZ, gabazine, GABA_A_ receptor blocker; TTX, tetrodotoxin voltage-gated sodium channel blocker; NPA, nipecotic acid, pan GABA transporter blocker; B1, Best1.

### Best1-independent component of TTX-insensitive GABA release is not mediated by the reverse mode of GATs

Not all of the non-synaptic tonic components disappeared in Best1 KO mice. In the young condition, a considerable amount (∼75.4%) of non-synaptic tonic component disappeared in the absence of Best1 (TTX insen. young B1 WT, 8.16 ± 0.92 pA, n = 17; young B1 KO, 2.01 ± 0.58 pA, n = 8). However, a significantly larger non-synaptic component remained in adult Best1 KO (Fig. 3d), indicating a non-synaptic and Best1-independent component. For this remaining component, we hypothesized that the reverse mode of GATs could be activated to compensate for the effect of Best1 KO in adult mice. To test this hypothesis, we measured TTX-sensitive and-insensitive tonic current under the blockade of GATs in the absence of Best1 (Fig. 4a-c). However, the blockade of GATs in the Best1 KO condition did not reduce any of the tonic GABA components (Fig. 4e), indicating no compensatory effect by the reverse mode of GATs.

**Figure 4.**
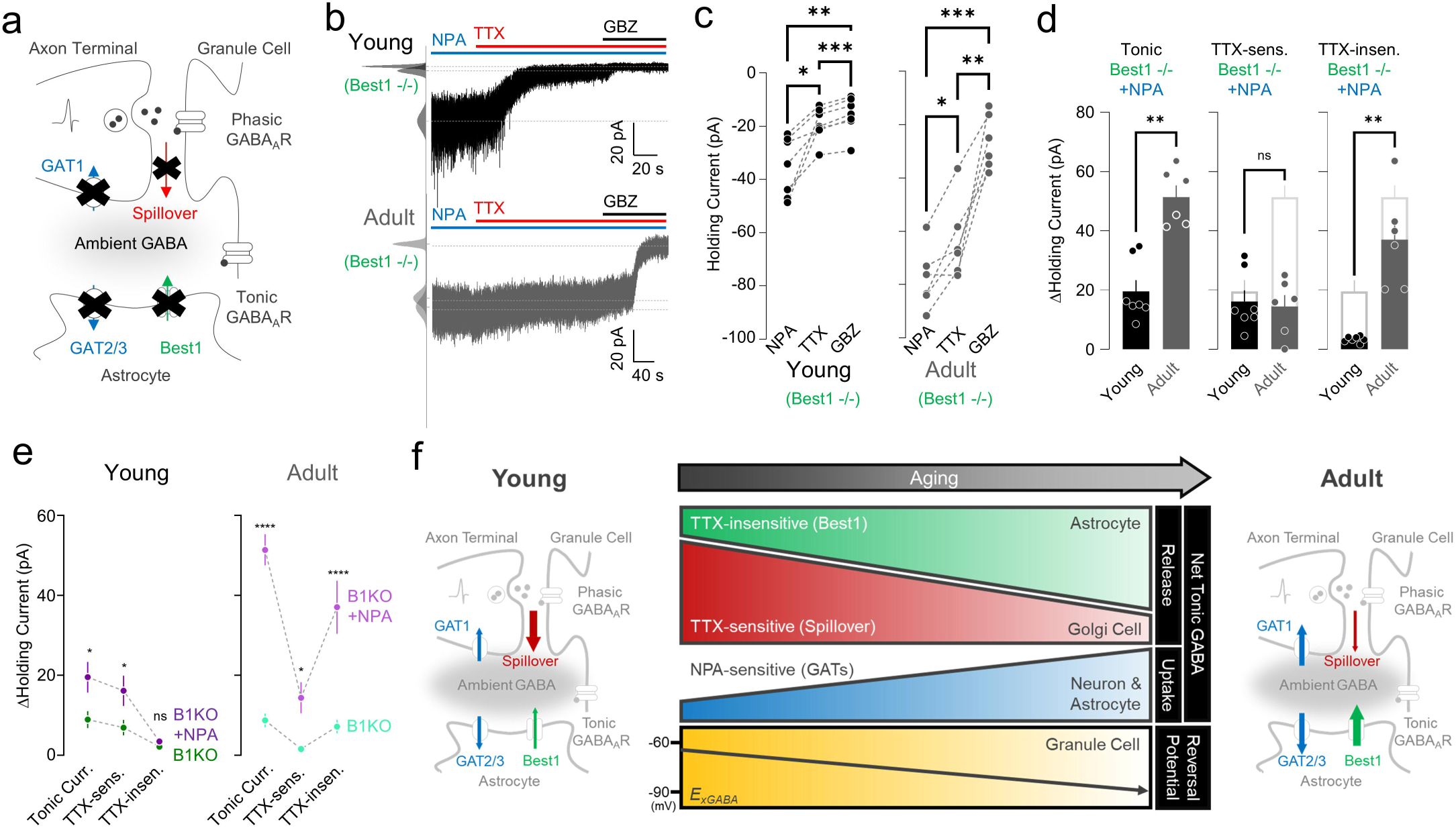
TTX-sensitive and TTX-insensitive tonic currents under blockade of GATs in absence of Best1. **a,** Model of GAT (blue), Best1-(green) and spillover-mediated (red) ambient GABA. **b,** Representative traces of GC tonic current from a young (top) and adult (bottom) mouse applied with TTX and GBZ under NPA presence in Best1-/-mice. **c,** Paired scatter plots of holding current change in young (left) and adult (right). Young: RM one-way ANOVA (**, p=0.0031), Tukey’s multiple comparisons test, NPA vs TTX, *p = 0.0125; NPA vs GBZ, **p = 0.0055; TTX vs GBZ, ***p = 0.0003; Adult: RM one-way ANOVA (***, p=0.0002), Tukey’s multiple comparisons test, NPA vs TTX (*, p=0.0336); NPA vs GBZ (***, p=0.0001); TTX vs GBZ (**, p=0.0058). **d,** Summary scatter bar graphs of tonic current (left), TTX-sensitive current (middle), and TTX-insensitive current (right). Mann Whitney test, Tonic: Young vs Adult, **p = 0.0012; TTX-sensitive: Young vs Adult, p > 0.9999; TTX-insensitive: Young vs Adult, **p = 0.0012. **e,** Comparison of B1 KO and B1 KO + NPA in terms of ΔHolding current in young (left) and adult (right) mice. Young: Two-way ANOVA, Sidak’s multiple comparisons test, Tonic: B1 KO vs B1 KO + NPA, *p = 0.0106; TTX-sensitive: B1 KO vs B1 KO + NPA, *p = 0.0309; TTX-insensitive: B1 KO vs B1 KO + NPA, p = 0.9734; Adult: Two-way ANOVA, Sidak’s multiple comparisons test, Tonic: B1 KO vs B1 KO + NPA, ****p < 0.0001; TTX-sensitive: B1 KO vs B1 KO + NPA, *p = 0.0230; TTX-insensitive: B1 KO vs B1 KO + NPA, ****p < 0.0001. **f,** Summary illustration of molecular and cellular switching of electrophysiological properties for tonic inhibition. GBZ: gabazine, GABA_A_ receptor blocker; TTX: tetrodotoxin voltage-gated sodium channel blocker; NPA: nipecotic acid, pan GABA transporter blocker.

**Figure 5.**
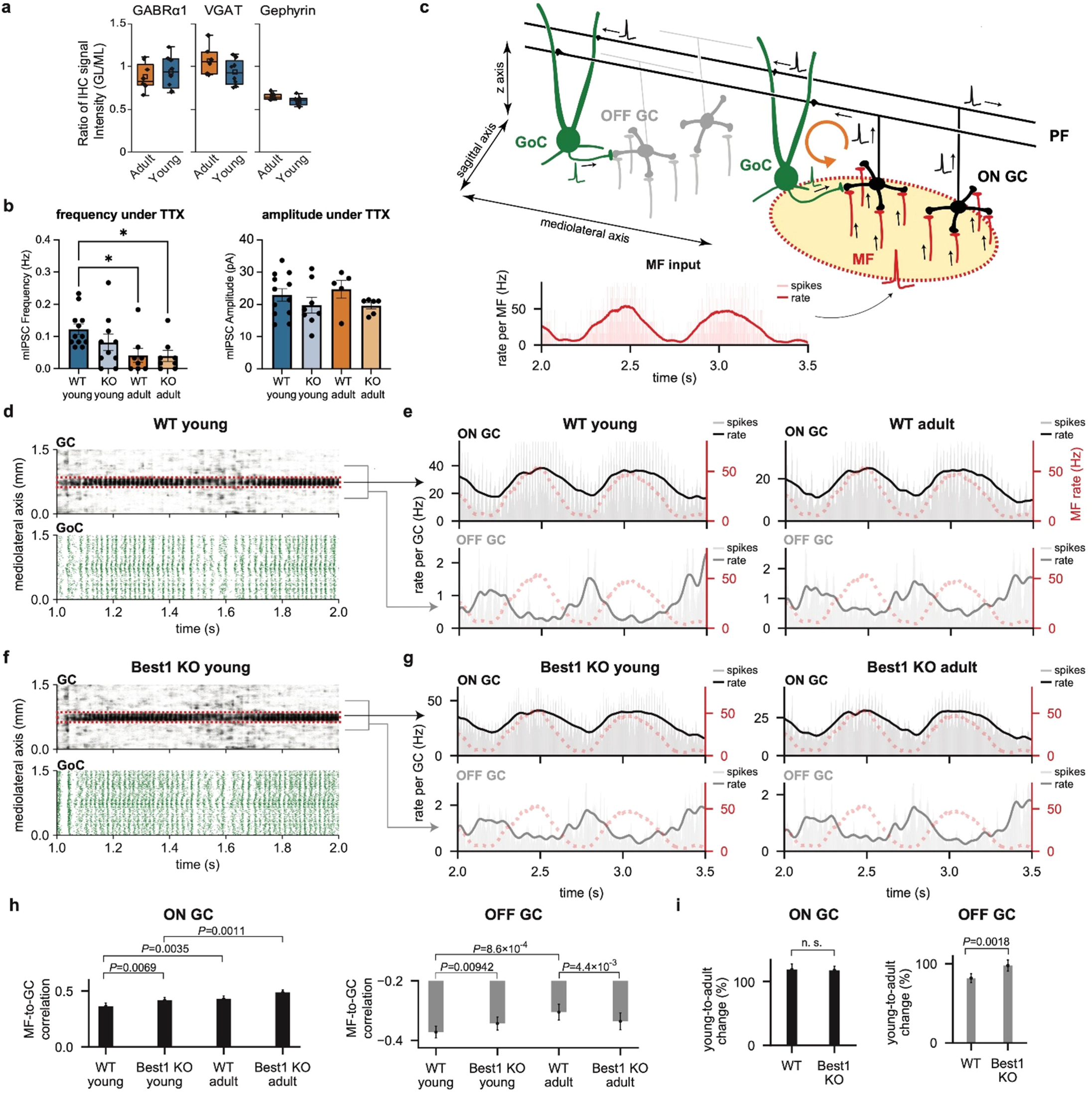
Network dynamics of *in silico* models with age-dependent tonic inhibition. **a**, Ratios of IHC signal intensities in GL relative to those in ML. **b**, mIPSC frequency and amplitude measured from young and adult animals in the WT and Best1 KO conditions. **c,** Network simulation protocol. In an excitatory input zone (yellow with a red border), MFs (red) transmit rate-modulated spiking activity (inset). GCs receiving these inputs (ON GC, black) project to GoCs (green) via PFs. GoCs in the zone form a feedback loop (orange arrow) to ON GCs while other GoCs inhibit off-zone GCs (OFF GC, gray). Inset: Spike histogram (light red bars, bin size=1 ms) and estimated firing rate (red line) of the input MFs. Rate was computed by gaussian filtering (*σ*=20 ms). **d,** Example raster plots of GC and GoC firings in the WT young animal-like condition. Spikes (dots) are aligned in the mediolateral axis. Red boxes mark ON GCs. **e,** Spike histograms (light-colored bars) and estimated rates (line) of ON GCs (Top, black) and OFF GCs (Bottom, grey). Red dotted lines show the MF rate in **a**. **f, g,** Same as b and c, but for the Best1 KO animal-like models. **h,** Correlation between MF and ON (Left)/OFF (Right) GC firings in each condition. **i,** Changes in correlations from the young-to adult-like condition. P-values are from the two-tailed t-test. Data are mean ± SEM (**c-g**) or 95% CI (**h**, **i**).

**Figure 6.**
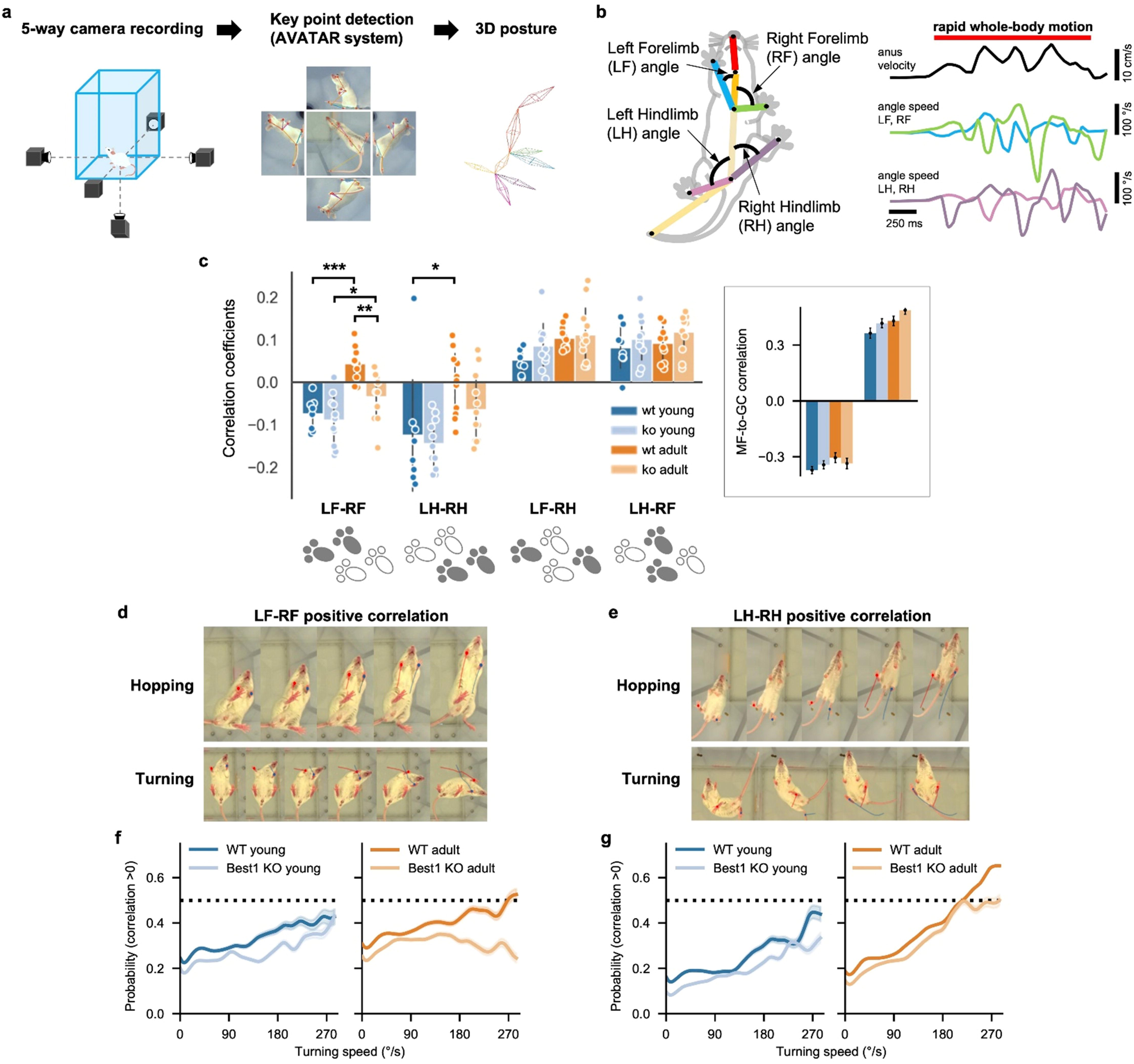
Impact of astrocytic GC tonic inhibition on the development of flexible movement coordination. **a,** Extraction process of 3D action skeleton using the AVATAR system. **b,** Left: Measured limb angles. Right: Example velocity of the anus node (top) and angular speeds of limbs (middle and bottom). A red line (top) denotes a period of rapid whole-body motion selected by anus velocity > 2 cm/s. **c,** Correlation coefficients of angular speeds between different limbs during rapid whole-body motion. *p<0.05, **p<0.01, ***p<0.001 (Two-way ANOVA and Tukey post-hoc test). Inset: MF-to-GC correlations in Fig. 5f, plotted in the same scheme for comparison. **d, e,** Example movements with positive forelimb (d) and hindlimb (e) correlations. **f,** Probability of positive forelimb correlation with respect to turning speed in the young (Left) and adult (Right) animals. **g,** Same as f but for hindlimb correlation. Data are mean ± SEM.

In the absence of both Best1 channels and GAT activity, we found that the amounts of TTX-sensitive components are similar between young and adult conditions (Fig. 4d). In contrast, we observed noticeably differential effects of GATs on the TTX-insensitive components: NPA uncovered minimal TTX-insensitive component in the young, whereas NPA uncovered significantly higher TTX-insensitive component in the adult (Fig. 4e), suggesting the emergence of an unknown, non-synaptic, Best1-independent component in adults, which requires future investigations.

### The reversal potential of extra-synaptic GABA_A_R, E_xGABA_, is more hyperpolarized in the adult GCs

So far, we have presented the developmental changes primarily in terms of tonic inhibitory current. Given that tonic inhibition provides substantial shunting conductance in GCs^12^, it is also crucial to check the age dependence of the inhibitory reversal potential of extra-synaptic GABA_A_R (E_xGABA_). Due to the technical challenges associated with obtaining E_xGABA_ of GCs using perforated patch voltage-clamp measurements, we adopted an alternative approach using THIP (4,5,6,7-tetrahydroisoxazolo[5,4c]pyridin-3-ol, a.k.a Gaboxadol)^21^, a selective agonist for delta-subunit-containing extrasynaptic GABA_A_ receptors.

As previously described^21–24^, we conducted cell-attached recordings of extrasynaptic GABA_A_R currents and voltage-gated K^+^ currents to assess the driving force for chloride ions in the presence of THIP (DF_THIP_) and the membrane voltage (V_m_), respectively (Supplementary fig. 1a,b). While the resting membrane potential V_m_ showed no significant difference between young and adult groups (Supplementary fig. 1e,f), DF_THIP_ was significantly higher in the young condition (Supplementary fig. 1c,d). As a result, the estimated reversal potential, E_xGABA_, was significantly more hyperpolarized in adult GCs (Supplementary fig. 1g). Therefore, while the total amount of tonic current is similar in young and adult animals, the shunting effect of extrasynaptic GABA on GCs should be greater and more strongly inhibitory in adults than in young GCs. Fig. 4f summarizes our findings on the molecular and cellular developmental switch that modulates the ambient GABA and tonic inhibition in young and adult mice.

### Computational modeling predicts age-dependent changes in network-level input coding

Beyond the level of individual neurons, how would an age-dependent shift in the origin of tonic GABA impact network-level phenomena? Addressing this question in experiments would require monitoring the activity of many GCs *in vivo*, which is extremely difficult to date. We adopted an *in silico* approach: We re-tuned the large-scale, physiologically detailed model of the granular layer network^25^ based on our experimental data for different conditions and simulated the model to see how the network encodes external inputs (Fig. 5).

First, we investigated which age-dependent changes in the network can explain the developments in tonic GABA. For this, we built a model of GC inhibitory synapses based on the sIPSC data and estimated the condition where we can replicate the experimentally measured level of tonic inhibition (Fig. 1g,h; see Methods and Supplementary methods for details). In the adult case, both in the Best1 WT and KO condition, we reproduced the experimental data when one GC was inhibited by eight Golgi interneurons (GoC) on average, comparable to previous studies^25,34^. However, the young animal data required the four and three-times more GoC inputs for the WT and Best1 KO groups, respectively.

One explanation could be the pruning of inhibitory synapses^35^ in GCs during maturation. To examine this possibility, we compared the expression of key inhibitory synaptic molecules—the α1 subunit of the GABA receptor (GABRα1), vesicular GABA transporter (VGAT), and gephyrin—between adult (8–10 weeks old) and young (3–4 weeks old) mice. Immunohistochemical (IHC) staining patterns in both the granular (GL) and molecular layer (ML) appeared similar across the two age groups (Supplementary Fig. 3a–c). Quantitative analysis confirmed that normalized GL/ML signal intensity ratios did not differ significantly between adult and young mice (P = 0.637 for GABRα1; P = 0.104 for VGAT; P = 0.083 for gephyrin; Mann-Whitney test; n=8 mice each for adult and young) (Fig. 5a). These results suggest that inhibitory GoC-GC connections remain stable during this developmental period.

Another explanation can be the changes in the intrinsic properties of the GoC-GC connections. In particular, we analyzed miniature inhibitory postsynaptic currents (mIPSCs) under TTX in our data, which revealed a decrease of the mIPSC frequency to one-third and a half during maturation in the WT and Best1 KO animals, respectively (Fig. 5b). Therefore, we developed probabilistically activating, stochastic GoC-GC inhibitory synapse models and used them in the computational network model of the cerebellar granular layer^25,26^ to examine how the differences in the probabilistic synaptic transmission and their contribution to GC tonic inhibition impact the granular layer network activity across different conditions.

The model simulation results demonstrated significant disparities in network activity between the young and adult-like WT conditions caused by differences in the activity-dependent (synaptic and spillover tonic) and independent (non-spillover tonic) inhibition. We simulated the network model with physiologically realistic, localized mossy fiber (MF) inputs with slowly oscillating firing rate modulation^25,36,37^ (Fig. 5c). Those inputs gave rise to increased activity in a cluster of GCs^25,38^ directly receiving the MF input (ON GCs) while the rest (OFF GCs) received inhibition mediated by parallel fibers (PF) and GoCs (Fig. 5d-g). In all cases, The firing rates of ON GCs faithfully followed those of the MF inputs, slightly better in adults. However, the anti-phasic modulation of OFF GCs, driven by the long-range GoC inhibition, was significantly weaker in adults, particularly in the WT condition (Fig. 5e,g).

Examining the age-dependent changes, we found that the activity of ON GCs became equally better correlated to MF input in adults, whereas OFF GCs became significantly less anti-phasic to MFs with age, much more in the WT than Best1 KO condition (Fig. 5h). This phenomenon was related to differences in activation of the excitation-inhibition loop of ON GCs and GoCs (orange arrow in Fig. 5c), which generates fast oscillations in the network activity^25,39^. When we compared the power of the MF input-driven and oscillatory components in the ON GC activity between the cases, the strength of the input-driven component remained stable across the conditions, whereas the network-generated oscillation component significantly decreased with age in the WT but not in the Best1 KO condition (Supplementary fig. 4). Since GoCs fire synchronously with the network-generated oscillation^25,39,40^, the weakened oscillation in the WT-like network in adults explained the decrease of inhibition-driven patterning of OFF GC firings in the same condition.

In summary, our simulations suggested that the shift from activity-dependent to independent inhibition, occurring from young to adult animals, can significantly decrease the PF and GoC-mediated mutual inhibition between GCs clusters innervated by different MF inputs, thereby promoting independence in their activity. In the context of motor coordination, if the MF inputs into distinct zones convey efference copies of upstream movement commands for different body parts, the stronger neuronal activity-independent (i.e., weaker activity-driven) inhibition in the adult granular layer can facilitate more flexible coordination of those body parts. This prediction was examined at the behavioral level in the next section.

### Shift in the origin of tonic inhibition enables flexible coordination of spontaneous movements

To test our prediction from the *in silico* models, we investigated how animals in each condition coordinate their movements using AVATAR, a system for monitoring animal behavior by high-speed cameras and extracting the posture online^27^ (Fig. 6a). This approach allowed us to assess motor coordination during a diverse array of spontaneous movements in an uncontrolled setting without any fixed behavioral tasks. We focused particularly on the coordination between limbs during whole-body motion, where the cerebellum plays a crucial role^41^. We first characterized the movements of individual limbs by calculating how quickly their relative angles to the trunk changed in time (angular speeds), followed by evaluating the correlations between the angular speeds across different limbs^29,30^ during rapid whole-body motion defined by the speed of an anus marker exceeding 2 cm/s (Fig. 6b). We anticipated finding largely negative correlations between opposing (left and right) fore-or hindlimbs since they should move in opposite directions during walking and running.

On the contrary, we found that the left-right correlations were significantly shifted toward a positive direction in adults, much more in the WT condition (Fig. 6c). To exclude whether this is due to simultaneous raising of the forelimbs during rearing, we re-analyzed data with an additional condition of forelimbs being no higher than the vertical positions of hindlimbs (see Methods), only to find the same pattern (Supplementary fig. 5). Finally, we divided the posture time-series data into distinct segments based on similarities in the multi-scale movement features (Supplementary fig. 6a; see also Methods) and again found significantly more segments with positive correlations in the WT adult animals (Supplementary fig. 6b).

Then, why did the positive left-right correlations arise when those animals moved around? A closer examination of animal behavior revealed a variety of motions with both sides of the fore-or hindlimbs moving concurrently, particularly in the WT adults (Fig. 6d, e). One example is hopping, where animals jump by supporting themselves with both left and right limbs together. Another example is twist turning, which involves swinging both fore-or hindlimbs in the same direction. However, despite a few easily recognizable examples, we noted that those movements mostly appeared in varying degrees, often subtly, rather than making isolatable behavior types. For instance, we found that the probability of positive correlation monotonically increased with the average turning speed for each motion segment, which is significantly less pronounced in the Best1 KO adults (Fig. 6f,g and Supplementary fig. 7). Therefore, animals gradually employed moving both sides of limbs together as their motion became more agile whereas the Best1 KO adults did not.

Furthermore, we also performed analysis of our 3D kinematic dataset using classical cerebellar function metrics^42^ (stride variability, ataxia index, inter-limb phase variance; Supplementary fig. 8). This revealed that Best1-KO mice do not exhibit increased variability; if anything, they show a trend toward reduced stride-time and step-width variability, consistent with greater locomotor stability. In contrast, our flexibility-specific analyses also identified clear deficits in adaptive inter-limb coordination, indicating that Best1 deficiency biases locomotion toward stereotyped, stable patterns, thereby impairing stability–flexibility trade-off (see more in Discussion).

In summary, our behavioral data and analysis supported the hypothesis raised by our computational models that the age-dependent shift from activity-dependent to independent inhibition in the cerebellar granular layer network promotes more flexible coordination of the movements of individual limbs. Therefore, it suggests that the shift in the origin of inhibition toward the astrocyte-released GABA is a crucial mechanism for developing a broad repertoire of whole-body motion from young to adult animals.

## Discussion

In this study, we have explored the developmental change in the sources of tonic inhibition on cerebellar GCs and its implications in neural network computation and behavior. For this, we combined three different experimental and computational methods of *ex vivo* characterization of cellular properties, *in silico* modeling based on experimental data, and finally, *in vivo* monitoring and analysis of spontaneous animal behavior aided by deep learning technologies. Our multi-faceted approach has allowed for a comprehensive understanding of how molecular and cellular properties impact neural network computation and, subsequently, behavior.

Our study is the first to describe the molecular and cellular developmental switch of tonic GABA inhibition during adolescence and toward adulthood in the mouse cerebellum (Fig. 4f). Our *ex vivo* characterization highlights that although the overall ambient GABA levels remain unchanged from youth to adulthood, the detailed molecular and cellular mechanisms of GABA release and uptake evolve with age. Initially, tonic GABA mainly originated from neuronal spillovers triggered by synaptic activities in young mice. As they age, astrocytic release through Best1 becomes the dominant source. Simultaneously, GABA uptake by GATs increases in both neurons and astrocytes. These developmental changes in GABA release and uptake contribute to the developmental switch of tonic GABA inhibition. In addition, the observed shift in E_xGABA_ of GC from-65 mV in young mice to-80 mV in adult mice also contributes to the developmental switch, providing stronger tonic inhibition in adults. These findings (Fig. 4f) shed light on the intricate inner workings of tonic GABA signaling within the GC layer, providing a detailed understanding that serves as a foundation for realistic modeling of information processing in the cerebellum.

Our study contests a widely accepted notion that, in mice, the cerebellar cells and circuit reach maturity by the age of three to four weeks^43^ before the onset of puberty, suggesting the same for cerebellum-dependent behavior. We demonstrated that tonic inhibition, a crucial regulator of GC excitability^8,18,44,45^, still undergoes a substantial transition after three weeks^7,14,16^ as described above. We also demonstrated that this transition parallels late-stage development in motor behavior, allowing for diverse and flexible movement coordination. This observation confirmed the prediction of our computational modeling that the changes in tonic inhibition in individual GCs can make network-level computation more flexible for encoding diverse inputs. While our study focused more on the role of GoC-independent inhibition, the importance of GoC-dependent inhibition is indisputable^10,46,47^. Although GC inhibition by individual GoCs is known to be weak and sparse^46,48,49^, the network dynamics of GoCs, such as highly correlated/synchronized activity^25,39,40,50^ driven by a GrC-GoC loop, is necessary. The powerful modulation of GrC activity by GoCs^47^ suggests such a collective activity of GoCs to perform adaptive thresholding on GrCs. However, population imaging data revealed that the activity of GoCs is highly correlated while it still is high-dimensional rather than in complete synchrony^50^. Our computational modeling, combined with experimental data, showed that a balance between the activity-dependent (synaptic, adaptive) and independent (extra-synaptic, non-adaptive) thresholding is closely related to that between collectivity versus independence in the Golgi cell population activity. We further showed that this balance follows the developmental trajectory where the activity-independent inhibition growing with age diminishes the network-wide impact of synchronized GoC-dependent synaptic and spillover inhibition, allowing more independent variabilities, or dimensionality, in the network activity^50–52^. Therefore, our work provides a novel insight about how the synaptic and extra-synaptic inhibition contribute together to make adaptive thresholding powerful enough but also suited for high-dimensional computation^50–52^.

Our analysis of spontaneous animal movements aligns with the network dynamics in our computational models but seemingly contradicts previous studies that showed apparently normal or even superior performance in motor tasks with impaired cerebellar tonic inhibition^18,45^. However, adaptive motor coordination can still be impaired even when movements appear normal. For instance, children with typical development can walk and run proficiently by the age of 10 years, but their locomotor adaptation is significantly slower than adults^19^, suggesting that more cognitive aspects of motor behavior, such as adaptation and flexible control, continue to develop beyond childhood^19,20^. Based on our results, we suggest that astrocyte-driven tonic inhibition plays a crucial role in that process.

Although we identified astrocytic Best1 as a main component of ambient GABA in adults, we also found undetermined sources beyond synaptic spillover, astrocytic Best1 channels, and GATs. One candidate is taurine, an agonist of various types of GABA_A_Rs, including the *δ* subunit-containing extrasynaptic GABA_A_Rs^53,54^. Given a taurine-positive signal in cerebellar Purkinje cells of adults^17^, Purkinje cells may tonically release taurine to tonically inhibit GCs, which is an intriguing possibility for future investigation. Our study supports that tonic inhibition evolves into a complex phenomenon during maturation, involving increasingly more diverse mechanisms^7^. Another mechanism to compensate for decreased tonic inhibition is homeostatic plasticity increasing the outward K^+^ current^55^. This mechanism can offset the changes in the baseline excitability but cannot remove all the effects due to decreased tonic inhibition. For example, the rheobase of the granule cells is the same between the WT and Best1 KO animals^18^ (Supplementary fig. 2) while the gain of the f-I curve is greater in the Best1 KO, indicating that the homeostatic compensation cannot fully replace tonic inhibition. We incorporated this effect (Supplementary fig. 2) into our simulations, which still exhibited the differences in the network activity. Therefore, homeostatic plasticity and tonic inhibition are not only replacements for each other but can play slightly different roles.

The seemingly contradictory reports of enhanced rotarod performance in Best1-KO mice^18^ versus impaired flexible coordination in our study can be reconciled by considering locomotor control as comprising two interacting dimensions—stability and flexibility. For example, reduced stride-time and step-width variability of Best1-KO mice (Supplementary fig. 8) are consistent with greater locomotor stability and in line with previous observation of improved rotarod performance^18^. On the other hand, clear deficits in adaptive inter-limb coordination suggest stereotyped, stably patterned movements that are advantageous in repetitive tasks like rotarod while it is detrimental for flexible motor adjustments. This stability–flexibility trade-off framework, supported by age-dependent rotarod performance^56^ and the inter-limb anti-correlation principle^29^, provides a mechanistic explanation for how Best1 loss can simultaneously enhance one aspect of motor behavior while impairing another, thereby confirming and extending the existing literature (see also Supplementary fig. 9 for summary).

One limitation of the present study is the use of constitutive Best1 knockout mice, which inherently complicates interpretation. The lifelong absence of Best1 may engage compensatory mechanisms that alter cerebellar circuit organization and function. Consequently, the behavioral phenotypes we report may reflect not only a disruption of the proposed adolescent developmental switch, but also earlier developmental abnormalities and/or ongoing compensatory changes throughout life. While acute and stage-specific manipulations of Best1 would be required to disentangle these possibilities, such approaches were beyond the scope of the current work. We acknowledge this limitation while noting that the convergent evidence from our *ex vivo*, *in silico*, and *in vivo* analyses consistently supports a critical role for Best1-mediated tonic inhibition in the maturation of motor coordination.

Our observation of the drastic changes in tonic inhibition throughout adolescence coincides with similar observations made in different brain regions, such as the hippocampus^57,58^, underscoring the importance of ambient GABA and related mechanisms in this developmental period. Therefore, we suggest that the development of tonic inhibition, driven by astrocytic mechanisms, is crucial in orchestrating the maturation of neural circuits and behavior from youth to adulthood.

## Acknowledgments

This work is supported by the Institute for Basic Science (IBS), Center for Cognition and Sociality (IBS-R001-D2) funded by the Ministry of Science to C.J.L. and S.H.; Okinawa Institute of Science and Technology Graduate University to E.D.S., S.H. and the NRF grant funded by the Korea Ministry of Science and ICT (RS-2024-00338607) to K. T. Y..

## Data and code availability

The source data and computer codes for simulations and analysis will be provided upon acceptance.

## Declarations

The authors declare no conflict of interest.

## Supplementary information

1. Supplementary methods
2. Supplementary figure 1-9
3. Supplementary table 1
4. Supplementary movie 1-4

## Notes

### Competing Interest Statement

The authors have declared no competing interest.

### Summary of Updates

All figures were revised; computational model has been updated based on experimental results from histology and eletrophysiology; Supplemental files were updated

