## Supplementary information for "Cerebellar tonic inhibition orchestrates the maturation of information processing and motor coordination"

### Supplementary methods

#### Construction and simulation of the granular network model

We built and simulated the computational models of the granular layer, based on a previously published model[1]. Briefly, the model simulated the granular layer of the cerebellar cortex with dimensions of 1.5 mm (mediolateral axis) × 0.7 mm (sagittal axis). It contained about 0.8 million granule cells (GC), 2000 Golgi cells (GoC), and 2000 mossy fibers (MF). Except MFs that delivered the external inputs, all the cells were based on active membrane conductance and intracellular mechanisms determined by experimental data[2]. The GC model had a single compartment while the GoC model had a somatic, two basal and two apical dendritic compartments. We computed the connectivity between the cells by the Peter's rule[3] determined either by the connection probability versus distance data in published studies and our data (see below) or by the collision between virtual axons and dendrites. In the latter case, we used our efficient implementation of the algorithm, Pycabnn[4].

#### Intrinsic properties of the cell models and their tuning

In this study, we made two-dimensional comparisons between young and adult animals and also between Best1-WT and KO groups. First, in the young versus adult case, we assumed that the cell intrinsic properties are already mature in young animals. Second, in the Best1-WT versus KO case, we assumed that Best1-KO triggers the changes in the subthreshold $K^+$ channels in GCs to maintain the baseline excitability. Simply removing the tonic inhibitory conductance caused the additive change in the GC f-I curve, similar to the one in pharmacological block experiments[5] (gray line in Supplementary fig. 2), while the Best1-KO case showed a multiplicative change with no shift in the rheobase[6]. To replicate this phenomenon, we tested various parameters changes in the $K^+$ channel models in the GC model and found that parameter changes in the inwardly rectifying (Kir) and M current-generating $K^+$ channels can induce the multiplicative change in the f-I curve with the level of tonic inhibition in our data (red line in Supplementary fig. 2). This GC model parameters were used as a baseline model in the KO case.

### Modeling of GC inhibitory synapses in GCs

To model individual inhibitory synapses in GCs, we first computed the auto-covariance function (ACF) of the sIPSC time series data from experiments. If an spontaneous synaptic event at t = t' is denoted by $sIPSC_1$, the sum of sIPSCs triggered by random synaptic events that occurred at {t1, t2, t3, · · · , tN } is

$$sIPSC(t) \;=\; \sum_{i=1}^{N} sIPSC_1(t-t_i) \;=\; \int_{-\infty}^{t} dt' p(t') sIPSC_1(t-t') \;=\; \int_{0}^{\infty} ds\, p(t-s) sIPSC_1(s)$$

where $p(t) \;= \sum_{i=1}^{N} \delta(t-t_i)$. Then,

$$ACF(\tau) = \langle sIPSC(t) sIPSC(t+\tau)\rangle_t \;=\; \mathcal{N} \iiint dtdsds' p(t-s)p(t+\tau-s') sIPSC_1(s) sIPSC_1(s')$$

where $\mathcal{N}$ is a normalization constant. Here we assume that synaptic events occurred randomly as in a Poisson process, which leads to

$$ACF(\tau) \approx \mathcal{N} \iiint dtdsds'(s-s'+\tau) sIPSC_1(s) sIPSC(s') \approx \mathcal{N}'\, ACF_1(\tau) \qquad \text{(S1)}$$

where $ACF_1(\tau) = \langle sIPSC(t) sIPSC(t+\tau)\rangle_t$. $\mathcal{N}'$ is also a normalization constant. Therefore, the ACF of sIPSC is the same as ACF of $sIPSC_1$ up to a constant factor. Furthermore, this will still be approximately true if sIPSC is caused by multiple synapses as long as the temporal profiles of $sIPSC_1$ caused by them are close to each other.

We assumed that $sIPSC_1$ has a double exponential form,

$$sIPSC_1(t) = \alpha_1 exp(-(t-t')/\tau_1) + \alpha_2 exp(-(t-t')/\tau_2) \qquad \text{(S2)}$$

where $\tau_1 < \tau_2$. We evaluated an analytic expression of $ACF_1$ from Eq. S2 and compared the result with sIPSC-ACF computed from the experimental recordings. In this way, we obtained model parameters ($\tau_i$, $\alpha_i$) for each recording for the WT-adult, WT-young, KO-adult, and KO-young cases and took average values to build computational synapse models ($sIPSC_1$). We also tested a three exponential model, but it did not retrieve another meaningful component.

### Modeling of tonic inhibition and GoC-to-GC connectivity

We investigated the conditions that can replicate the level of tonic inhibitory current observed in the voltage clamp experiments by simulation of the models. To do so, we inserted the inhibitory synapse model that we built (see above) in the GC model and ran the simulation of the voltage clamp experiment with the holding potential of -60 mV at room temperature. The spontaneous inhibitory events at each synapse were modeled as a random Poisson process occurring at 3 Hz rate.

The maximum synaptic conductance parameters were set to the value computed from experimental data. We compared two different measurements in experiments, the peak sIPSC and reversal potentials of $K^+$ and $Cl^-$ ions, and estimated the synaptic conductance. Since the estimates were comparable between Best1 WT and KO animals if they belonged to the same age group, we used the average across the WT and KO animals for both cases.

Finally, we set the tonic conductance of Cl− ions in the cell model to the value calculated from the experimental data for the TTX-independent tonic IPSC. In the adult Best1 WT and KO cases, those parameters could reproduce the level of tonic IPSC in experimental data, with eight GoC inputs in the original network model[1]. However, we found that the number of GoC inputs had to be 24 and 16 in the young Best1 WT and KO cases, respectively, while experimental data showed that this is entirely attributed to the differences in probabilistic synaptic transmission (Fig. 5a-b). Therefore, we used the GoC-to-GC convergence of 24:1 for the simulations in all conditions. On the other hand, we developed a stochastic synapse model in NEURON that transmits presynaptic spikes with a probability $P_{release}$. We used $P_{release}$ = 1, 0.333, 0.666, and 0.333 for the WT young, WT adult, Best1 KO young, and Best1 KO adult conditions, respectively.

### Construction and simulation of the network models

After determining all relevant parameters of the network models (see Supplementary Table 1), we built and simulated them on the OIST cluster computer Deigo (https://groups.oist.jp/scs/deigo), following the previously described procedure[1,4]: We first determined the cell positions in the models, generated the virtual dendrites and axons, and computed the connectivity between the cells via our custom package, Pycabnn[4]. We also prepared the spike times of external MF inputs based on the stimulation protocols[1]. Then, we ran the simulation of the models via the NEURON 8.2 simulation platform[7].

We first ran the baseline simulation with MFs randomly firing at 5 Hz with adjusting a bias current in model GCs until they fired at ~0.5 Hz. We also simultaneously adjusted the conductance of GC-GoC synapses until GoCs fired at ~3 Hz in average, resulting in 200 pS for all cases. After determining all the parameters using this procedure, we proceeded to simulate the network with different external MF patterns.

### Segmentation, clustering, and analysis of paw movement time series

We segmented the paw movement time series of each mouse in the following way: We first up-sampled the time series to 100 Hz and collected the movement features at each time point (Fig. 6a). The first set of the features was from the spectrograms of paw velocities in an ego-centric frame[8] where an anus and chest node are fixed by translation and rotation of all marker positions. The spectrograms were computed by wavelet transformation in the frequency range from 0.1 Hz to 10 Hz by a step of 0.1 Hz. Then, we applied logarithms and took 20 principal components, which explained more than 99% of the total variance. The second set of the features included the speed of the anus and chest nodes in the horizontal plane, body height defined by the first principal components of the vertical marker positions, and rate of change in the body height. We reduced the dimensionality of the feature vectors down to two dimensions by the t-SNE algorithm (perplexity: 50)[9]. This procedure separated the movement time series into segments with the similar movement features. We clustered the segments by the hierarchical density-based clustering algorithm[10] with the Euclidean distance and the threshold of 1, and selected the large movement clusters where the anus velocity is greater than 2 cm/s and the forelimb-to-hindlimb vertical distance is less than 2.576 times the standard deviation of the hindlimb-to-hindlimb vertical distance.

For each large movement cluster, we computed the cross-correlation of the angular speeds for the left and right forelimbs and hindlimbs (Fig. 6b). We also computed the rotation speed by computing the horizontal angle of the anus-to-chest vector and taking its time-derivative. The probability of positive correlation versus body rotation speed (Fig. 6f,g) was computed in the following way: We first evaluated a density of the body rotation speed ω in all the large movement clusters, P(ω), and another density of ω only in those with positive (LF-RF or LH-RH) correlations, P(ω|+), by kernel density estimation. Then, by Bayes' theorem,

$$P(+|\omega) = \frac{P(\omega|+)P(+)}{P(\omega)}$$

where P(+) is the fraction of the large movement clusters with positive correlations. We also used the procedure of repeating the density evaluation 1000 times with randomly selecting the large movement clusters with replacement to compute the mean and SEM of P(+|ω).

All velocities were computed by the Savitzky-Golay filter (window size: 4 ms, order: 3). Spectrograms were computed by using PyWavelets package[11]. Dimensionality reduction and clustering were performed by Scikit-learn package[12] .

### Supplementary figures

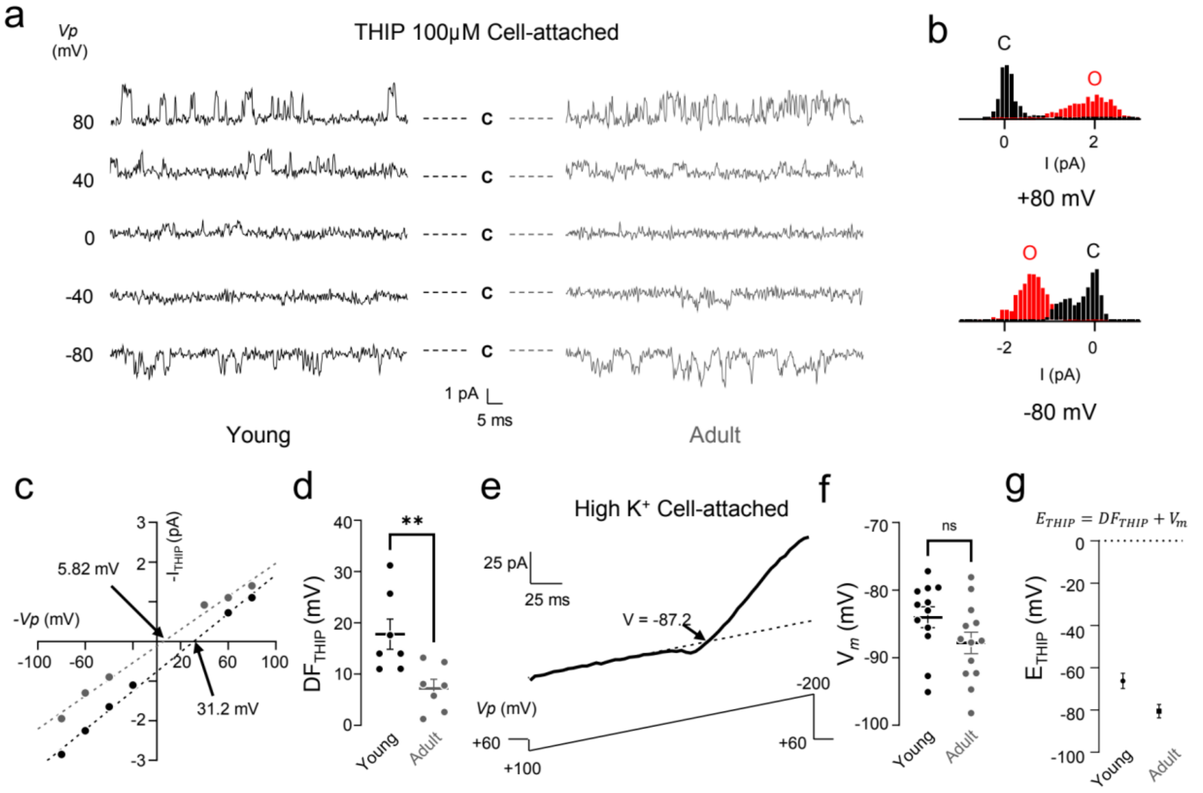


**Supplementary figure 1** Reversal potential of extra-synaptic $GABA_AR$ is higher in young compared to adult mice. **a**, Single channel recording with a cell-attached patch with THIP, selective δ-subunit containing extra-synaptic GABAAR agonist. **b**, Representative histograms from channel recording trace at +80 mV (top) and -80 mV (bottom). **c**, Representative IV curve linearly fitted for detection of THIP-mediated driving force. **d**, Summary scatter plot of THIP mediated driving force in young (17.79 ± 2.94 mV) and adult (7.28 ± 1.71 mV) mice. Unpaired Student's t-test, Young vs. Adult, **p = 0.0093. **e**, Representative trace of high potassium cell-attached patch with ramp protocol. Unpaired Student's t-test, Young vs. Adult, p = 0.1035. **f**, Summary scatter plot of membrane potential in young (-87.81 ± 1.60 mV) and adult (-84.02 ± 1.55 mV) mice. **g**, Summarized point plot of calculated reversal potential of extra-synaptic GABAAR between young (-66.23 ± 3.62 mV) and adult (-80.53 ± 3.21 mV) mice. Unpaired Student's t-test, Young vs. Adult, p < 0.0001. THIP: Gaboxadol.

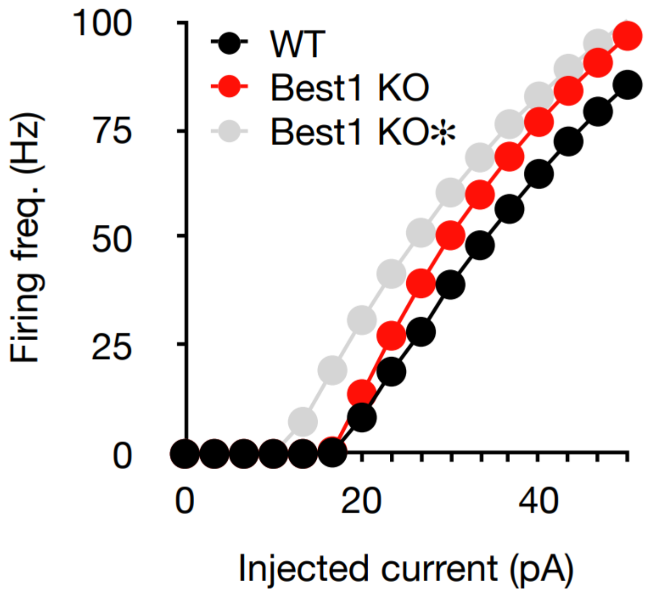


**Supplementary figure 2** Firing rate versus injected current (f-I) curve of the model GCs. Black: Baseline GC model for the Best1-WT case. Red: Model with decreased tonic inhibitory conductance and adjusted parameters in the low-threshold $K^+$ channels for the same rheobase as WT. Gray: Model only with decreased tonic inhibitory conductance.

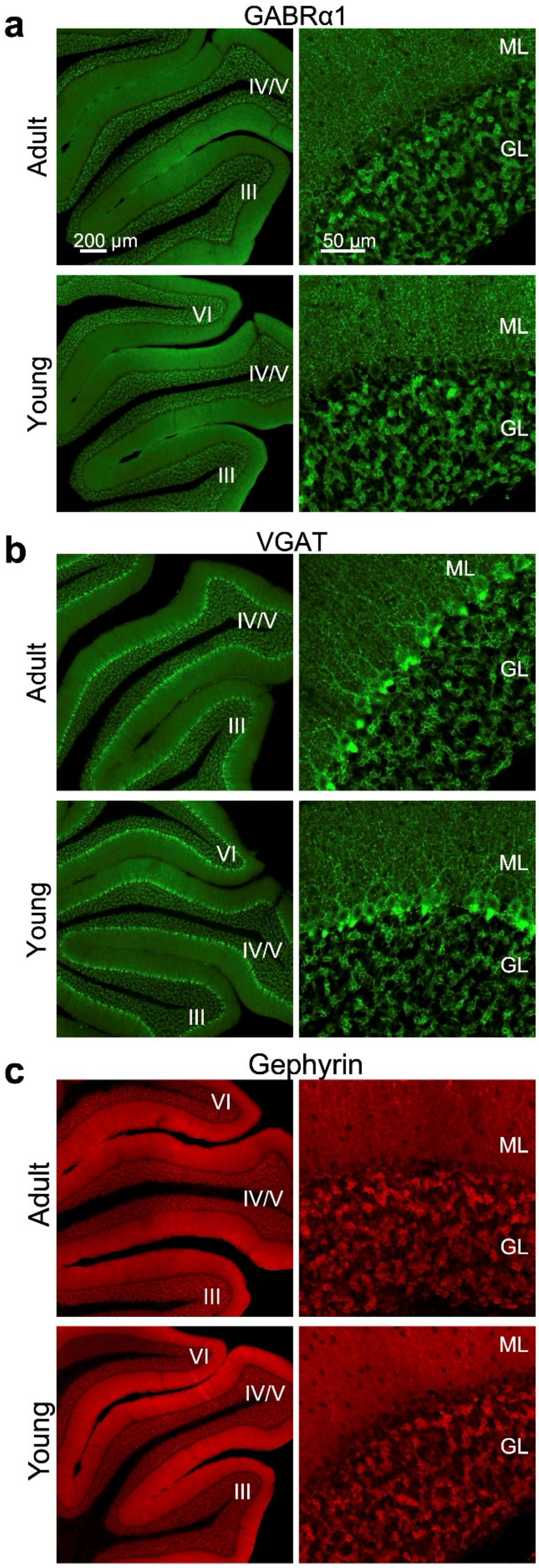


**Supplementary figure 3** Representative images of adult (upper) and young (lower) cerebellar slices stained with antibodies against GABRα1 (**a**), VGAT (**b**) and gephyrin (**c**). The roman numbers in low-magnification images indicate the lobules.

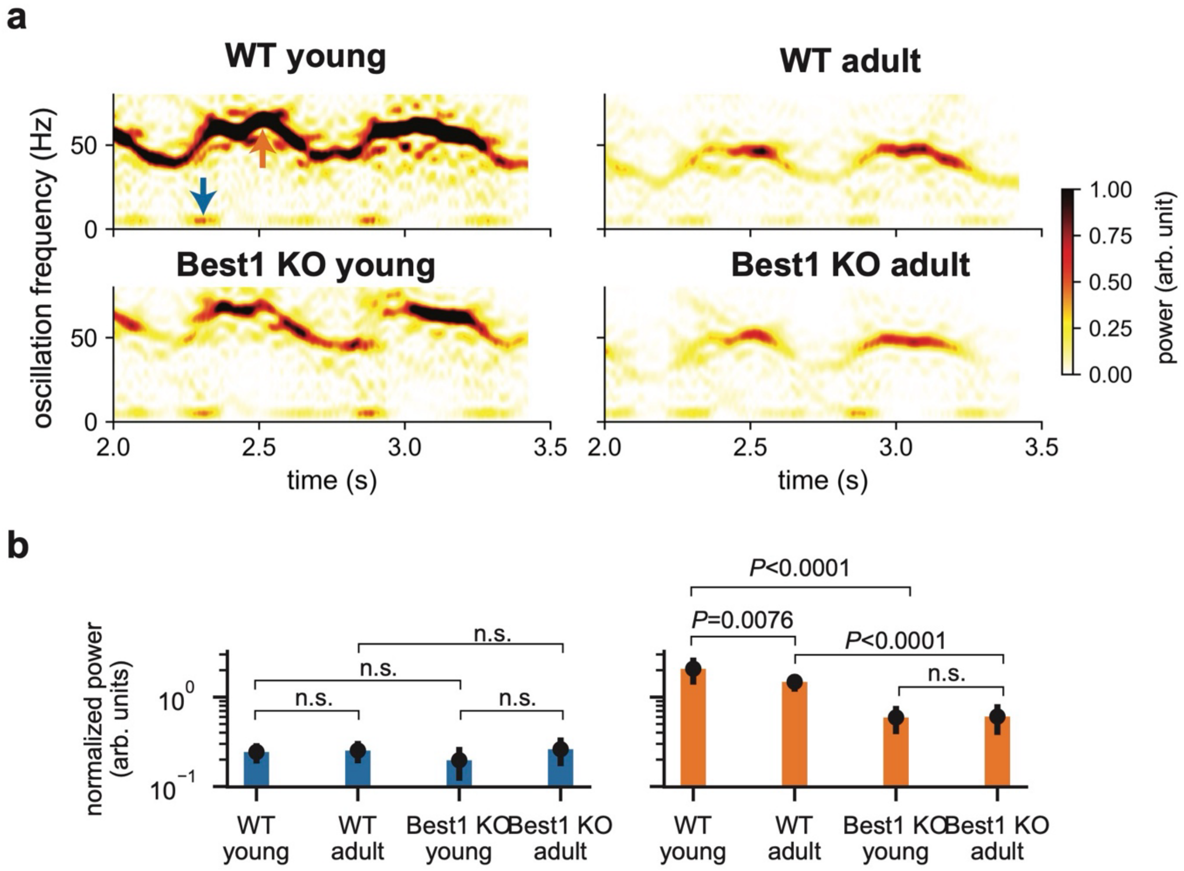


**Supplementary figure 4** Input-driven versus network-generated GC activity in each condition. **a**, Spectrograms of the spike histograms in Fig. 5e, g. Arrows point to the MF input-driven low- (blue) and network-generated high-frequency (orange) oscillations, analyzed in **b**. **b**, Left: Power of the low frequency, MF input-driven oscillations in GC firings (blue in **a**) normalized by the mean firing rate. Right: Same for the high frequency, network-generated oscillations (orange in **a**). Data are mean ± SEM. P-values are from the two-tailed t-test. n.s.: P > 0.05.

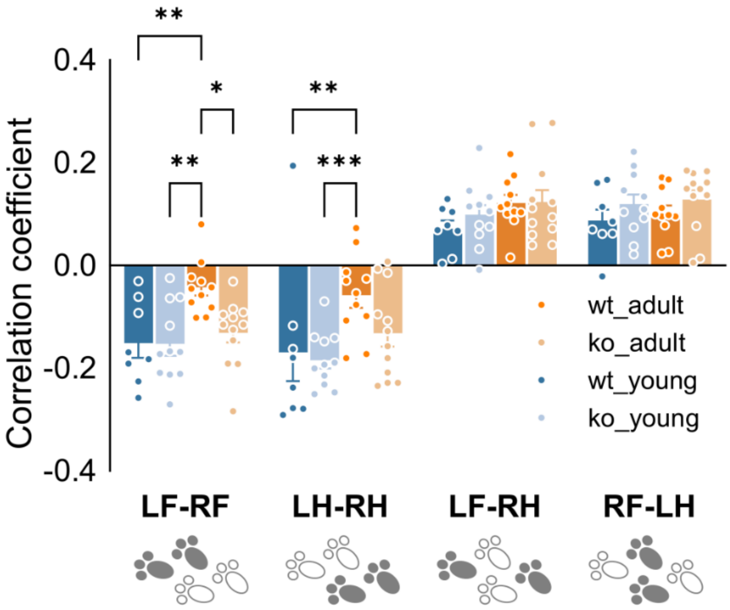


**Supplementary figure 5** Correlation coefficients of angular velocities between forelimbs and hindlimbs during rapid movements as in Fig. 6c, with an additional condition that forepaw elevation is not higher than the hindpaw level. *p<0.05, **p<0.01, ***p<0.001 (Two-way ANOVA and Tukey post-hoc test).

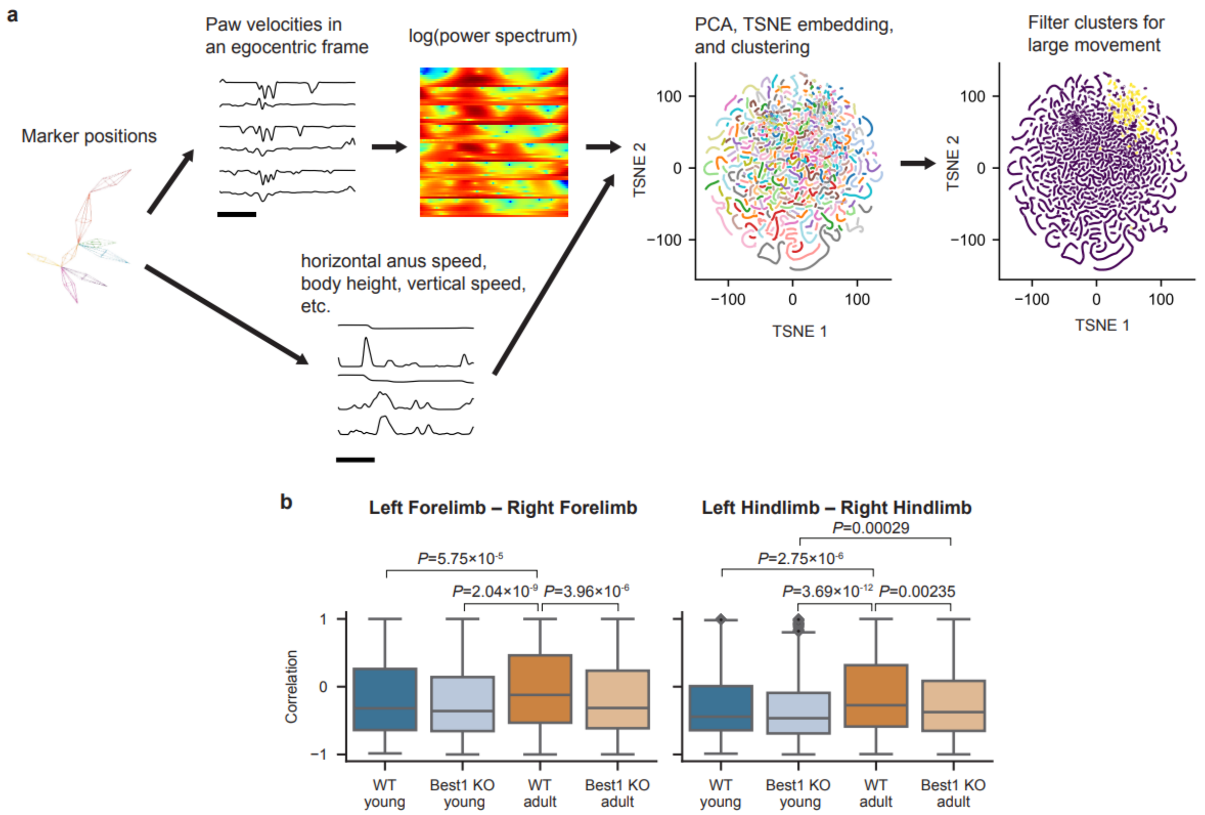


**Supplementary figure 6** Analysis of paw movement time series by using feature-based segmentation and clustering. **a**, Schematic diagram of the segmentation and clustering procedure. **b**, Cross-correlation of the angular speeds of the left and right forelimbs (Left) and hindlimbs (Right) in the rapid motion clusters (selected by the same condition as in Supplementary fig. 4; yellow in a) for each category. P-values are from the Komogorov-Smirnov test.

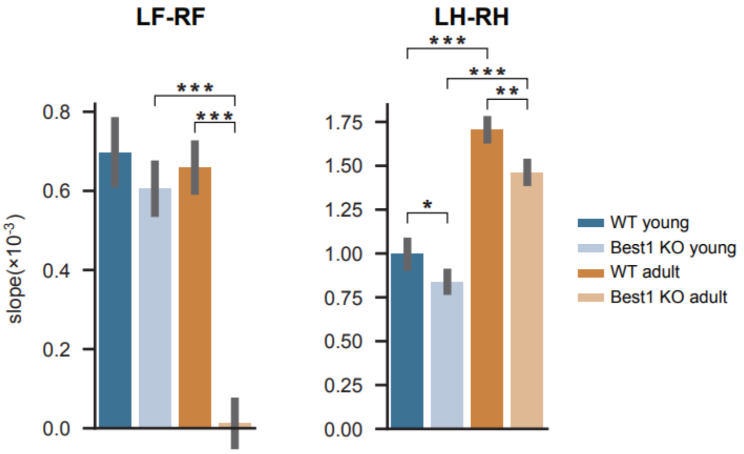


**Supplementary figure 7** Slopes of the turning speed versus the probability of positive correlations in Fig. 6f,g. Comparisons are made only between the adult and young or between the Best1 WT and KO conditions. ***: P<0.001, **:P < 0.01, *:P < 0.05. P-values are from the two-tailed t-test. Data are mean ± SEM.

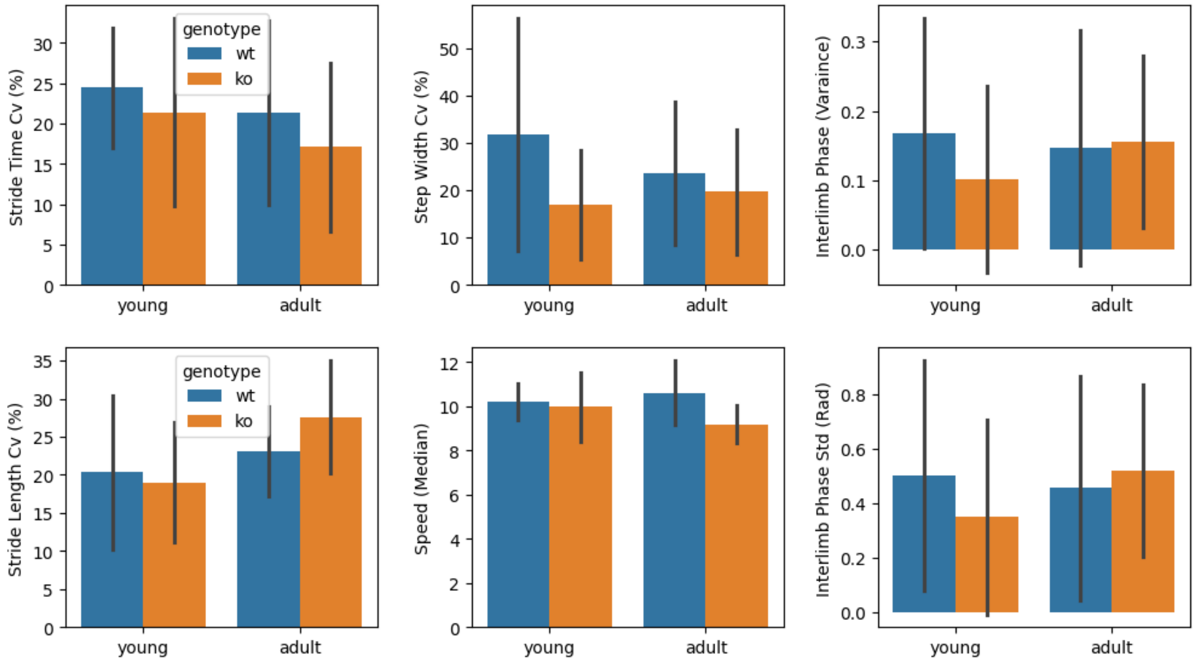


**Supplementary figure 8** Analysis of traditional stride-based cerebellar function metrics from 3D kinematic dataset. Conventional cerebellar function analyzed with 3D kinematic data using stride-based measures. Hind-paw steps were detected from smoothed velocity signals (20 Hz), with strides defined between consecutive left foot-strikes and right steps paired using a midpoint criterion to ensure robust alignment. For each stride, we quantified stride duration, stride length, step width, duty factors, and temporal symmetry. From these, we derived variability indices (CV% of stride time, stride length, step width) and inter-limb phase statistics (circular mean, variance, SD). Mean ± SD.

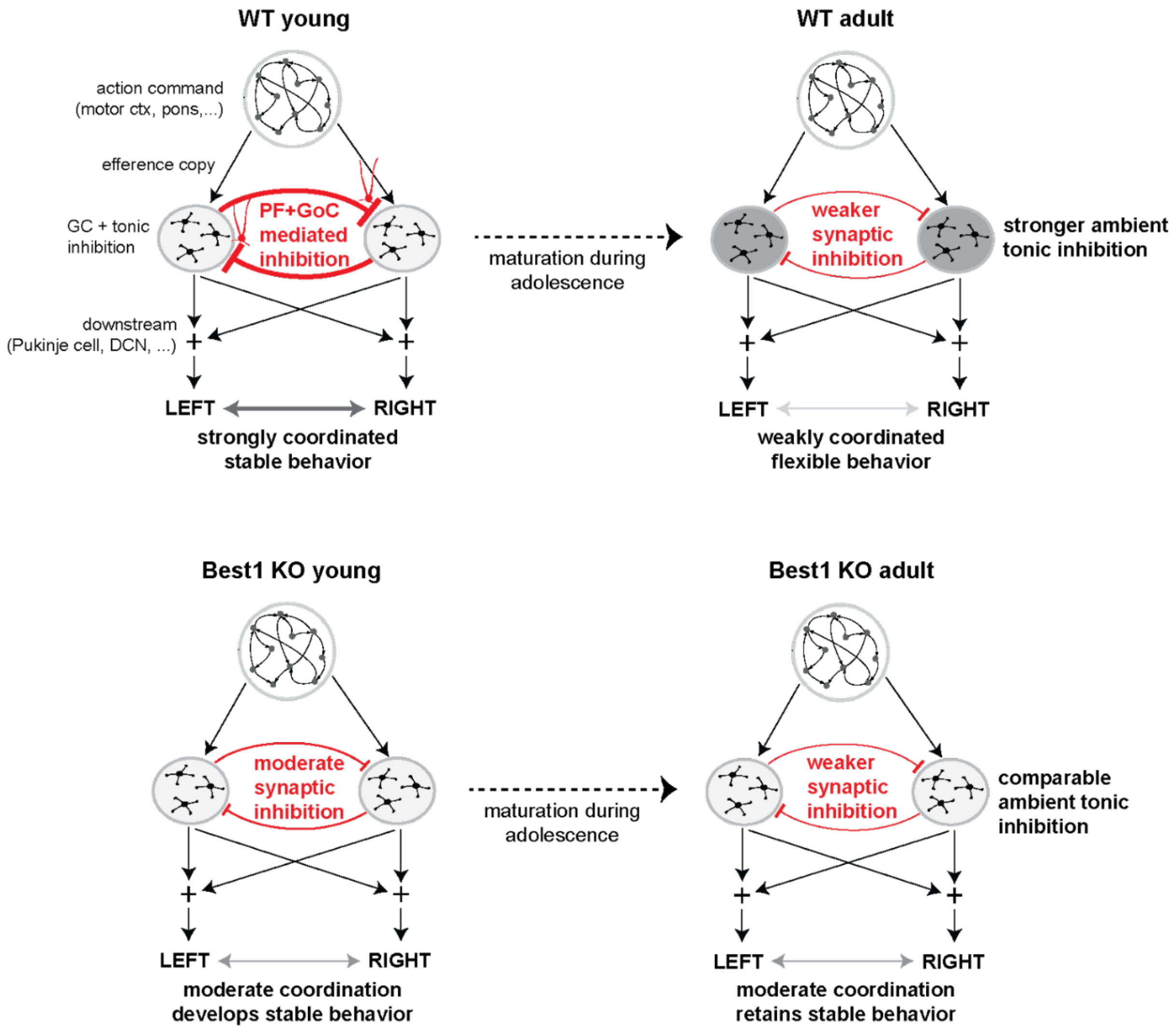


**Supplementary figure 9** Conceptual models for our results. **Top Left,** Simplified neural network model of the cerebellum-mediated movement coordination. Efference copies of neural action commands to muscles enter the cerebellum and innervate GCs in different microzones. They inhibit each other via GoCs activated by PFs projected from the other microzone. The GC activity in those microzones are combined to form the coordination outputs to modify the action commands. If the mutual inhibition is strong as in the WT young animals, this leads to strong movement coordination between the left and right side, which stabilizes movement. **Top Right,** The same as Top Left but for the WT adult case after maturation. The mutual inhibition significantly weakens while glia-mediated ambient tonic inhibition grows and maintains GC excitability. This leads to weaker movement coordination that allows more variable and flexible movements. **Bottom,** The same as Top but for Best1 KO animals. The mutual inhibition and ambient tonic inhibition much less significantly change during maturation than WTs. This leads to much less significant changes in behavior stability and flexibility.

### Supplementary tables

| Case | $\tau_1$ (ms) | $\tau_2$ (ms) | $g_{syn}$ (pS) | $E_{K+}$ (mV) | $E_{Cl-}$ (mV) | $g_{tonic}$ (μS/cm$^2$) |
|---|---|---|---|---|---|---|
| Best1-WT Adult | 9 | 81 | 520 | -88 | -80.53 | 116.4 |
| Best1-WT Young | 4.71 | 22.2 | 520 | -88 | -66.23 | 73.9 |
| Best1-KO Adult | 6.8 | 95 | 340 | -88 | -80.53 | 55.1 |
| Best1-KO Young | 4.5 | 33 | 340 | -88 | -66.23 | 15.7 |

**Supplementary table 1** Model parameters in each case. $\tau_1$ and $\tau_2$ are decay time constants of synaptic inhibitory conductance. $g_{syn}$ is a mean maximal conductance for GoC-to-GC inhibitory synapses. $ECl^-$ is the $Cl^-$ reversal potential. $g_{tonic}$ is the specific conductance of the tonic inhibitory ($Cl^-$) conductance. All other parameters are the same as the model in previous study[1].

### Supplementary movies

**Supplementary movie 1** Activity of neurons in our computational granular layer network model in the WT young condition. Black and green dots represent firing of GCs and GoCs, respectively. For visual clarity, each dot (spike) lasted for 5 ms from its spike time. A red circle represents the region with the rate-modulated MF inputs, depicted in Fig. 5c. The time range is also from 2 sec to 2.8 sec of the simulation, identical to Fig. 5c.

**Supplementary movie 2** Activity of neurons in the network model in the WT adult-like condition. The notations are the same as Supplementary movie 1.

**Supplementary movie 3** Activity of neurons in the network model in the Best1 KO young-like condition. The notations are the same as Supplementary movie 1.

**Supplementary movie 4** Activity of neurons in the network model in the Best1 KO adult-like condition. The notations are the same as Supplementary movie 1.
